# Evolution of a fuzzy ribonucleoprotein complex in viral assembly

**DOI:** 10.1101/2025.04.26.650775

**Authors:** Huaying Zhao, Tiansheng Li, Sergio A. Hassan, Ai Nguyen, Siddhartha A.K. Datta, Guofeng Zhang, Camden Trent, Agata M. Czaja, Di Wu, Maria A. Aronova, Kin Kui Lai, Grzegorz Piszczek, Richard D. Leapman, Jonathan W. Yewdell, Peter Schuck

**Affiliations:** Laboratory of Dynamics of Macromolecular Assembly, National Institute of Biomedical Imaging and Bioengineering, National Institutes of Health, Bethesda, MD 20892, USA; Cellular Biology Section, Laboratory of Viral Diseases, National Institute of Allergy and Infectious Diseases, National Institutes of Health, Bethesda, MD 20892, USA; Bioinformatics and Computational Biosciences Branch, National Institute of Allergy and Infectious Diseases, National Institutes of Health, Bethesda, MD 20892, USA; Electron Microscopy Unit, Trans-NIH Shared Resource on Biomedical Engineering and Physical Science, National Institute of Biomedical Imaging and Bioengineering, National Institutes of Health, Bethesda, MD 20892, USA; Biophysics Core Facility, National Heart, Lung, and Blood Institute, National Institutes of Health, Bethesda, MD 20892, USA; Laboratory of Cellular Imaging and Macromolecular Biophysics, National Institute of Biomedical Imaging and Bioengineering, National Institutes of Health, Bethesda, MD 20892, USA; HIV Dynamics and Replication Program, Center for Cancer Research, National Cancer Institute, Frederick, MD 21702, USA

**Author notes:** Correspondence to Peter Schuck.

## Abstract

Previously we showed that the genetic diversity of SARS-CoV-2 nucleocapsid (N) protein explores a wide range of biophysical properties facilitated by non-local impact of point mutations to its intrinsically disordered regions (Nguyen et al., 2024). This includes modulation of self-association, such as the creation of a *de novo* binding interface through the P13L mutation characteristic of Omicron variants. In the present work we focus on the key function of N condensing viral RNA into ribonucleoprotein particles (RNPs) for viral assembly. Lacking high-resolution structural information, biochemical and biophysical approaches have revealed architectural principles of RNPs, which involve cooperative interactions of several protein-protein and protein-RNA interfaces, initiated through oligomerization of conserved transient helices in the central disordered linker of N. Here we study the impact of defining N-protein mutations in variants of concern on RNP formation, using biophysical tools, a virus-like particle assay, and reverse genetics experiments. We find convergent evolution in repeated, independent introduction of amino acid substitutions strengthening existing binding interfaces, compensating for other substitutions that promote viral replication but decrease RNP stability. Furthermore, we show that the P13L mutation of Omicron variants enhances RNP assembly and increases viral fitness. Overall, our data reveal RNP complexes to be highly variable not only in sequence and conformations, but also in thermodynamic and kinetic stability, with its pleomorphism affecting basic architectural principles. We hypothesize that the formation of polydisperse, fuzzy N-RNA clusters with multiple distributed weak binding interfaces optimizes reversible RNA condensation, while supporting host adaptation and allowing for a large sequence space to be explored.

## Introduction

Intrinsically disordered proteins are ubiquitous and play key role in many dynamic cellular processes, including cellular signaling, transcriptional regulation, as well as spatio-temporal organization (Dyson and Wright, 2005; Holehouse and Kragelund, 2024). Intrinsic disorder is particularly prevalent in proteins of RNA viruses, for reasons deeply connected to evolution in presence of their high mutation rates and quasispecies nature (Brown et al., 2011; Gupta and Uversky, 2024; Tokuriki et al., 2009; Xue et al., 2014).

On a physicochemical level, disorder supports multi-functionality through modulation of the conformational ensemble dependent on ligands and post-translational modifications (Botova et al., 2024; Carlson et al., 2022, 2020; Jack et al., 2021; Ranganathan et al., 2023; Savastano et al., 2020; Syed et al., 2024) and the local environment (Nesmelova et al., 2019; Roden et al., 2022), the possibility of transient folding (Alderson et al., 2023; Bessa et al., 2022; Zachrdla et al., 2022; Zhao et al., 2022), and the propensity for protein clustering and liquid-liquid phase separation (LLPS) (Alberti et al., 2019; Brocca et al., 2020; Perdikari et al., 2020; Ranganathan and Shakhnovich, 2020; Savastano et al., 2020; Zachrdla et al., 2022). Disorder also permits a significant degree of sequence variation, as many functionally important features are not dependent on the specific sequence but are encoded in non-local biophysical properties (Alston et al., 2023; Nguyen et al., 2024; Zarin et al., 2021). Opposite to the achievement of mutational tolerance through high stability, such as exhibited, for example, by thermostable enzymes, mutational tolerance can be achieved in loosely packed and disordered proteins by lowering the potentially deleterious effect of mutations on stability (Tokuriki et al., 2009; Tokuriki and Tawfik, 2009). In addition, disorder and flexibility magnifies potential impact of mutations on conformation and biophysical properties (Nguyen et al., 2024). Furthermore, the high degree of sequence variability supports the exploration of short linear interaction motifs (SLiMs) in intrinsically disordered regions for interfacing with a large variety of eukaryotic regulatory and signaling processes, thus favoring adaptability and evolvability on the level of the host-virus interface (Davey et al., 2011; Duro et al., 2015; Hagai et al., 2014; Schuck and Zhao, 2023).

In recent years it has become apparent that assemblies of disordered proteins frequently retain significant disorder and conformational flexibility of their interaction partners, generating “fuzzy” complexes (Longhi et al., 2017; Tokuriki et al., 2009; Tompa and Fuxreiter, 2008). The potential functional advantages of intrinsic disorder in protein interactions include leveraging of weak binding interfaces through allovalency and the adaptability to multiple binding partners (Longhi et al., 2017; Olsen et al., 2017). In fuzzy complexes the total binding energy is distributed into multiple distinct ultra-weak interaction sites (Olsen et al., 2017). Similar to individual RNA virus proteins with loose or absent structure, maintaining disorder and a spatial distribution of low-energy interactions in the protein complexes may increase the tolerance for mutations and improve evolvability of protein complexes.

The unprecedented worldwide sequencing effort of SARS-CoV-2 genomes during its rapid evolution in humans provides a unique opportunity to examine these concepts. The genomic database now exceeds in size that of any other virus by orders of magnitude (Elbe and Buckland-Merrett, 2017; Rochman et al., 2022), providing the basis for phylogenetic analyses and for monitoring the emergence of variants of concern and their geographic spread (Hadfield et al., 2018). In addition, it exhaustively samples the mutational landscapes of amino acids that can occupy any position of the viral proteins, which reflects their biophysical constraints (Bloom and Neher, 2023; Nguyen et al., 2024; Zhao et al., 2022). For example, the SARS-CoV-2 nucleocapsid (N-)protein – the most abundant viral protein in infected cells and the focus of the present work – has 419 positions, 86% of which can be assumed by on average 4-5 different amino acids, and up to 12 in positions in the three intrinsically disordered regions (Zhao et al., 2022) (**Figure 1A**), evidently without fatally compromising dozens of reported N-protein functions (Wu et al., 2023). Observed N-protein mutations can modulate basic biophysical properties including its oligomeric state, thermodynamic stability of its two folded domains, LLPS propensity, charge distributions, and secondary structure content (Nguyen et al., 2024; Zhao et al., 2022), as well as its interactions with RNA (Cubuk et al., 2024; Dhamotharan et al., 2024), kinases (Johnson et al., 2022; Syed et al., 2024) and other host proteins through altered SLiMs (Li et al., 2025; Ren et al., 2024; Schuck and Zhao, 2023; Tugaeva et al., 2023).

**Figure 1.**
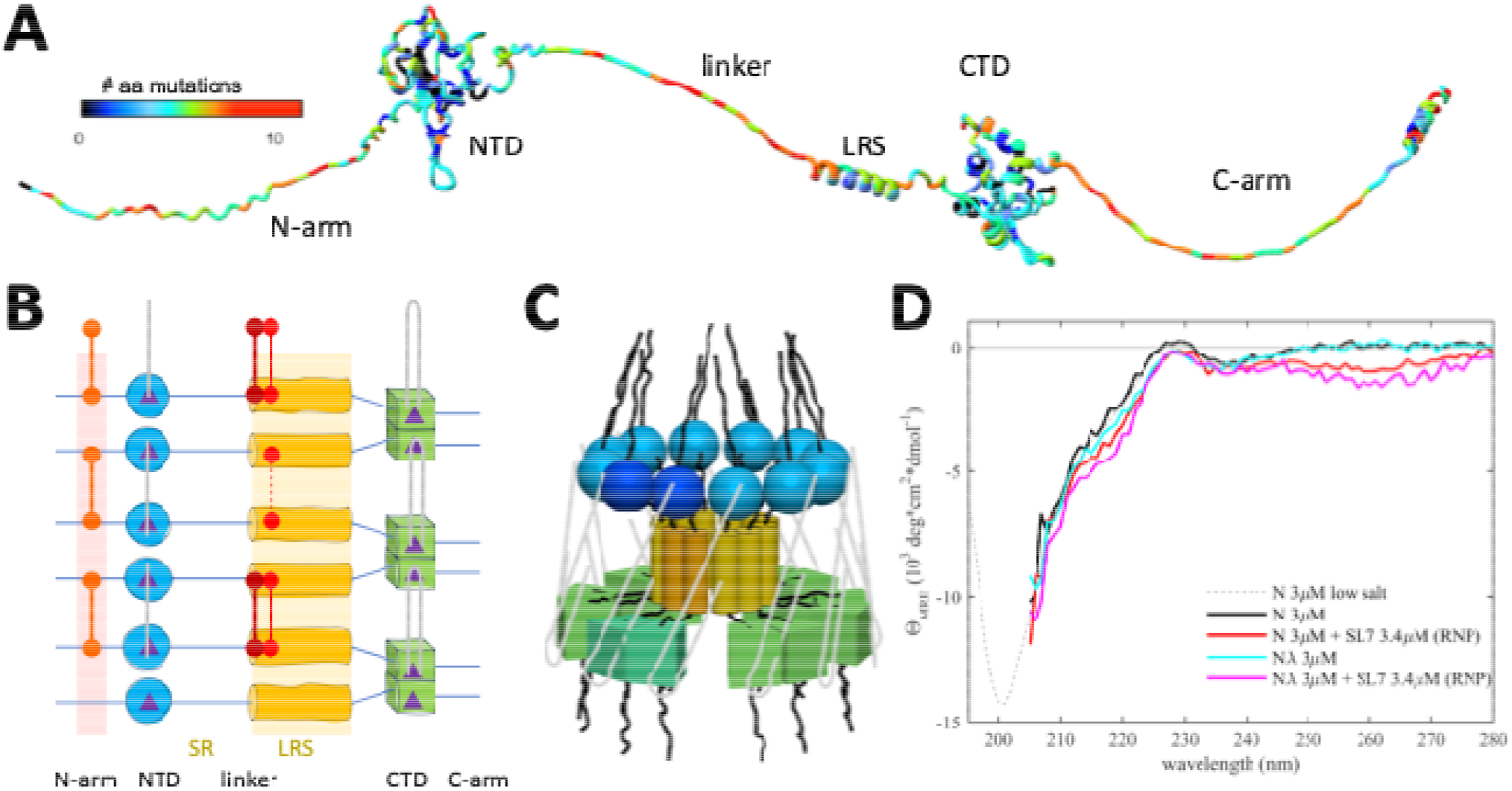
Basic organization of N-protein and RNPs. N-protein (1-419) has two folded domains, NTD (45-180) and CTD (248-363), and three intrinsically disordered regions including the N-arm (1-44), the central linker (181-247) and the C-arm (364-419). (A) Displayed is an AF2 structure where the disordered N-arm, linker, and C-arm are artificially stretched for clarity. The residues are color-coded according to the number of different amino acids that have been observed at this position in the mutational landscape replacing the Wuhan-Hu-1 sequence. The bioinformatic analysis was carried out as described, previously (Zhao et al., 2023), updated to August 26, 2024, using a threshold of >5 genomes for each mutation. (B) Schematic of protein-protein and protein/RNA interfaces in RNP assembly. The nucleic acid binding domain at the N-terminus (NTD) is indicated in blue, the LRS in yellow, and the dimerization domain (CTD) in green. Regions of self-association are indicated by shaded backgrounds. The linker is subdivided in a serine and arginine-rich region (180-205, SR) and a L-rich region (206-247, LRS). LRS can transiently fold into helices that create a hydrophobic patch for promiscuous self-association (indicated as yellow pattern). For clarity the cartoon only shows 3 neighboring N-protein dimers, although higher-order oligomers assemble in RNPs. Nucleic acid binding sites (purple triangles) preferentially bind single-stranded RNA at the NTD (grey lines), and double stranded RNA at the two sites per CTD dimer, with the ability to cross-link neighboring dimers potentially in various configurations. New inter-dimer interactions evolved in variants of concern are indicated by red connectors, including the promotion of beta-sheet oligomerization through the N:P13L mutation in the N-arm (as in Omicron and Lambda variants), and the introduction of cysteines at the base of the LRS helices in N:G214C (as in Lambda variants) and N:G215C (as in Delta variants). (C) Three-dimensional cartoon of the circular organization of N-protein domains in RNPs, with one dimer shaded slightly darker to highlight the dimeric building blocks. For clarity, subunit sizes are not drawn to scale. Alternate arrangements are depicted in **Supplementary Figure S1new.** (D) CD spectra of N-protein in the presence of SL7 under near physiological salt conditions leading to majority assembly of RNPs. Spectra are corrected for free SL7 contributions. Shown are spectra of ancestral N-protein alone (black), in the presence of SL7 forming RNPs (red), N_λ_ alone (cyan) and in the presence of SL7 forming RNPs (magenta). Spectra are truncated at < 205 nm due to limited buffer transparency. For comparison the dotted line shows a previously published CD spectrum of ancestral N-protein in low-salt buffer that permits measurement at shorter wavelengths (Nguyen et al., 2024). Triplicate scans yield average standard deviations of 0.13 (N), 0.17 (N+SL7), 0.16 (N_λ_), and 0.21 (N_λ_ +SL7) 103 deg cm2/dmol, respectively, with non-overlapping confidence bands for the different species, for example, between 215-220 nm.

In the present work we study the eponymous function of N-protein, which is the spatial condensation of the long genomic RNA (gRNA) into ribonucleoprotein particles (RNPs) for viral assembly. N-protein has two folded domains (the nucleic acid binding domain and the C-terminal dimerization domain, NTD and CTD, respectively), and three intrinsically disordered regions that include the linker between NTD and CTD, and the N-terminal and C-terminal extensions, N-arm and C-arm, respectively (**Figure 1A**) (Chang et al., 2006). The intrinsically disordered regions amount to ≈45% of total residues, which renders N-protein highly flexible with a radius of gyration fluctuating from 5 nm to >8 nm (Różycki and Boura, 2022). RNPs are ≈15 nm diameter particles composed of ≈10-15 N-proteins that bind stretches of gRNA (Klein et al., 2020; Yao et al., 2020). 30-40 RNPs are distributed like beads on a string in the ≈100 nm virion (Klein et al., 2020; Yao et al., 2020). RNPs appear highly heterogeneous in electron microscopy (EM) (Carlson et al., 2022; Landeras-Bueno et al., 2025; Yao et al., 2020), and as of now, a high-resolution structure has not been determined. The Morgan laboratory has described an *in vitro* model for the assembly of RNPs, where N-protein in the presence of stem-loops of the 5’-UTR RNA readily forms polymorphic ribonucleoprotein complexes that match in size, symmetry, and RNA content what would be expected for the assembled RNPs in virions (Carlson et al., 2022, 2020). In conjunction with biophysical experiments and point mutations exposing essential binding interfaces, this has allowed us recently to develop a coarse-grained structural model of RNPs (Zhao et al., 2024).To examine how architecture and energetics of RNP assemblies can be impacted by N-protein mutations we study a panel of N-proteins derived from ancestral Wuhan-Hu-1 and different variants of concern, including Alpha, Delta, Lambda, and Omicron (see **Table 1**), in biophysical experiments, VLP assays, and mutant virus. Specifically, we ask how the RNP size distribution and life-time is modulated by: (1) the novel binding interface created by the P13L mutation of Omicron; (2) enhancements of other weak self-association interfaces through G215C of Delta and G214C of Lambda; (3) the ubiquitous R203K/G204R double mutation of Alpha, Lambda, and Omicron. We also test whether the P13L mutation improves viral fitness, similar to G215C and R203K/G204R. The results are discussed in the framework of fuzzy complexes and molecular evolution of N in the course of viral adaptation to the human host. Understanding the salient features of the binding interfaces in viral assembly and their evolution expands our foundation for the design of therapeutics such as assembly inhibitors.

**Table 1.** Overview of N-protein mutant species studied.

| designation | N-protein mutations | in set of defining VOC mutations <sup>(1)</sup> | predominant oligomeric state at low μM concentrations | reaction boundary s-value in RNP assay (S) | MP final average Mw in 500-1500 kDa range (kDa) | best-fit mass increase from 0.3 to 3 μM (kDa) | best-fit effective RNP life-time (sec) |
| --- | --- | --- | --- | --- | --- | --- | --- |
| N (ancestral) | none | Wuhan-Hu-1 | dimer | 19.7 | 614 | 63.8 | 66.3 |
| N:P13L | P13L | $\lambda$, o (all) | | | | | |
| N:Δ31-33 | Δ31-33 | o (all) | | | | | |
| N:P13L/Δ31-33 | P13L, Δ31-33 | o (all) | dimer | 20.2 | 625 | 118.2 | 43.6 |
| N:R203K/G204R | R203K, G204R | $\alpha$, $\gamma$, $\lambda$, $\zeta$, and o (except XEC) | dimer | 19.5 | 596 | 70.3 | 54.6 |
| N<sub>o</sub> | P13L, Δ31-33, R203K, G204R | o (except XEC) | dimer | 20.5 | 617 | 77.2 | 58.5 |
| N:G215C | G215C (reduced) | $\delta$ (all 21J) | dimer/tetramer | 20.4 | 619 | 47.8 | 231 |
| N:G215C* | G215C (oxidized) | $\delta$ (all 21J) | tetramer | 20.8 | 649 | 56.1 | 41.8 |
| N<sub>λ</sub> | P13L, R203K, G204R, G214C (reduced) | $\lambda$ | dimer/tetramer | 21.0 | 669 | 20.0 | 67.4 |
| N<sub>λ</sub>* | P13L, R203K, G204R, G214C (oxidized) | $\lambda$ | tetramer | 21.5 | 660 | 51.6 | 98.2 |
| N:R203M | R203M | $\delta$, $\kappa$ | | | | | |
| N<sub>210-419</sub>* | Δ1-209 | o (not XEC) | | | | | |

<sup>(1)</sup> Referring to the most common mutations in the variants of concern, excluding sporadic spontaneous reversions or other variations.

## Results

### Essential protein-protein interfaces in RNP assembly

Based on biophysical experiments of protein-protein and protein-NA interfaces of full-length (FL) N-protein and domain constructs, in combination with the *in vitro* RNP assembly assay (Carlson et al., 2022, 2020) we have recently proposed a coarse-grained structural model of RNPs that satisfies all known binding interfaces. These include at least two known sites of protein-protein and two for protein-nucleic acid interactions distributed throughout most of the protein (Zhao et al., 2024) (**Figure 1B/C**). N-proteins are constitutive dimers with K_D_ in the low nM range by virtue of domain-swapped beta hairpins in the CTD (Chang et al., 2006; Cubuk et al., 2025; Yu et al., 2005; Zhao et al., 2021; Zinzula et al., 2021). N dimers exhibit only ultra-weak further self-association with ≈1 mM K_D_ (Yu et al., 2005; Zhao et al., 2022, 2021) (although nucleic acid impurities can artificially scaffold N-protein into higher oligomers (Carlson et al., 2020; Tarczewska et al., 2021)). However, higher-order assembly is initiated by the occupation of the nucleic acid binding site in the NTD, which allosterically promotes a helical conformation of the leucine-rich sequence (LRS) in the intrinsically disordered linker, allowing LRS helices to form promiscuous coiled-coil oligomers stabilized through hydrophobic interactions with low μM K_D_ (Zhao et al., 2023). As demonstrated with the LRS point mutant N:L222P that abrogates these transient helices, LRS oligomerization forms the basis for oligomerization of N-protein dimers into RNPs, for example, dodecamers as hexamers of dimers. These oligomers are stabilized by multi-site interactions with double-stranded RNA simultaneously in two sites of the CTD dimer, and by binding preferentially single-stranded RNA to each of the NTDs (Cubuk et al., 2024; Iserman et al., 2020; Korn et al., 2023; Roden et al., 2022), with both RNA interactions allowing inter-dimer interactions that crosslink N-protein dimers within the RNP (**Figure 1C**) (Zhao et al., 2024). Additional stabilizing protein-protein interfaces may exist in the C-arm (Carlson et al., 2022; Q. Ye et al., 2020), although we have been unable to detect self-association of the C-arm by analytical ultracentrifugation up to low mM concentrations (Zhao et al., 2023).

As suggested in the cartoon of **Figure 1C**, this supports the hypothesis of a three-dimensional arrangement with a central LRS oligomer with symmetry properties and dimensions similar to low resolution EM images of model RNPs (Carlson et al., 2022, 2020) and cryo-ET of RNPs in virions (Klein et al., 2020; Yao et al., 2020). It should be noted, however, that the arrangement sketched in **Figure 1C** is not unique and other subunit orientations could be envisioned that satisfy all constraints from experimentally observed binding interfaces, including different oligomers and anti-parallel subunits as illustrated in **Supplementary Figure S1**. Extending previous ColabFold structural predictions that show multiple N-protein dimers self-assembled *via the LRS coi*led-coils (Zhao et al., 2023), we attempted the AlphaFold modeling of RNPs combining multiple N dimers with SL7 RNA ligands, mimicking our biophysical assembly model. Current AlphaFold restrictions limit the prediction to pentamers of N-protein dimers with 10 copies of SL7 RNA. While only inconsistent results were obtained – which is not surprising given the large intrinsically disordered regions exceed the predictive power of AlphaFold – some models did produce an overall RNP organization similar to **Figure 1C**, suggesting such an arrangement is at least sterically reasonable with regard to possible N-protein subunit orientations in an RNP (**Supplementary Figure S2**).

Using the *in vitro* RNP assembly model of N-protein in mixtures with stem-loop RNA SL7 we measured secondary structure content in circular dichroism (CD) experiments (**Figure 1D**). While a significant structural change in the RNA when bound to N-protein can be deduced in the near UV, after correction for RNA contributions, in the far-UV the CD spectrum of the RNP mixtures is very similar to that of N-protein alone. A small gain in helicity can be discerned for RNP assembly mixtures at ≈220 nm, consistent with the expected coil transition in the LRS stabilizing the RNP. Similarly, small increases in helicity can be discerned for LRS helix-stabilizing cysteine mutants. It is noteworthy that the CD spectrum of the RNP mixture appears to be dominated by disordered chains. Although their signature minimum ellipticity at ≈200 nm cannot be observed directly in the high salt buffer required for RNP assembly, the data are quantitatively consistent in their strongly decreasing slope from 205 – 210 nm with the previously measured disordered signature of N-protein in low-salt buffer (without RNP formation). This result is consistent with the absence of significant structure formation in the RNP, and with N-protein intrinsically disordered regions retaining most of their disorder outside the LRS.

As is depicted in the diagram **Figure 1B**, assembly of approximately 6 N-protein dimers and ≈500bp of RNA requires simultaneous binding at multiple protein-protein and protein-RNA interfaces. Many of these interactions are weak, but they are multivalent and act cooperatively (Zhao et al., 2024). This feature may aid in dynamic assembly and disassembly, and shape the ensemble of complex states to efficiently populate functioning RNPs. As we describe in the following, mutations of N-protein have led to diverse mechanisms modulating and promoting RNP formation through their effect on protein-protein interfaces relevant to RNP stability.

### A novel self-association interface through transient β -sheets enabled by the P13L mutation

In early epidemiological analyses, the N:P13L mutation has been identified as the most important driver for N-protein fitness, and it has become an obligatory mutation of all Omicron variants (Obermeyer et al., 2022; Oulas et al., 2021). In our recent survey of biophysical effects of N-protein mutations relative to the ancestral protein, we observed a distinct ability of the N-arm mutant peptide N_1-43_:P13L to form large assemblies at ≈mM concentrations (which is not seen with the ancestral peptide), while full-length N:P13L exhibited enhancement of LLPS (Nguyen et al., 2024), both indicative of weak interactions. Furthermore, after prolonged storage of P13L N-arm peptide solutions at ≈mM concentrations at 4 °C, an increase in the solution viscosity was observed (Nguyen et al., 2024). In the present work we studied these effects of the P13L mutation in more detail. As shown in **Figure 2A**, negative-stained EM images show the formation of fibrils of N-arm peptides of Omicron variant N_1-43_:P13L,Δ31-33. Similarly, fibril formation was observed for N_1-43_:P13L N-arm peptide lacking the deletion Δ31-33, but not in controls with ancestral N-arm, the ancestral N-arm carrying only the deletion N_1-43_:Δ31-33, or the disordered ancestral C-arm (**Supplementary Figure S3**). Thus, the N-arm mutation P13L is responsible for the formation of fibrils in N-arm peptides after prolonged storage. Some of these N-arm fibrils exhibit a twisted morphology with width of ≈5 nm (**Figure 2A**), in some instances exhibiting patterns of strand breaks. Such fibrils are frequently encountered in proteins that can stack β-sheets, such as in amyloids (Paravastu et al., 2008). While we have not observed fibril formation in the context of full-length N, and have no evidence such fibrils are physiologically relevant, their occurrence in solutions of truncated N-arm peptide nonetheless demonstrates the introduction of ordered N-arm self-association interfaces in conformations of P13L mutants.

**Figure 2.**
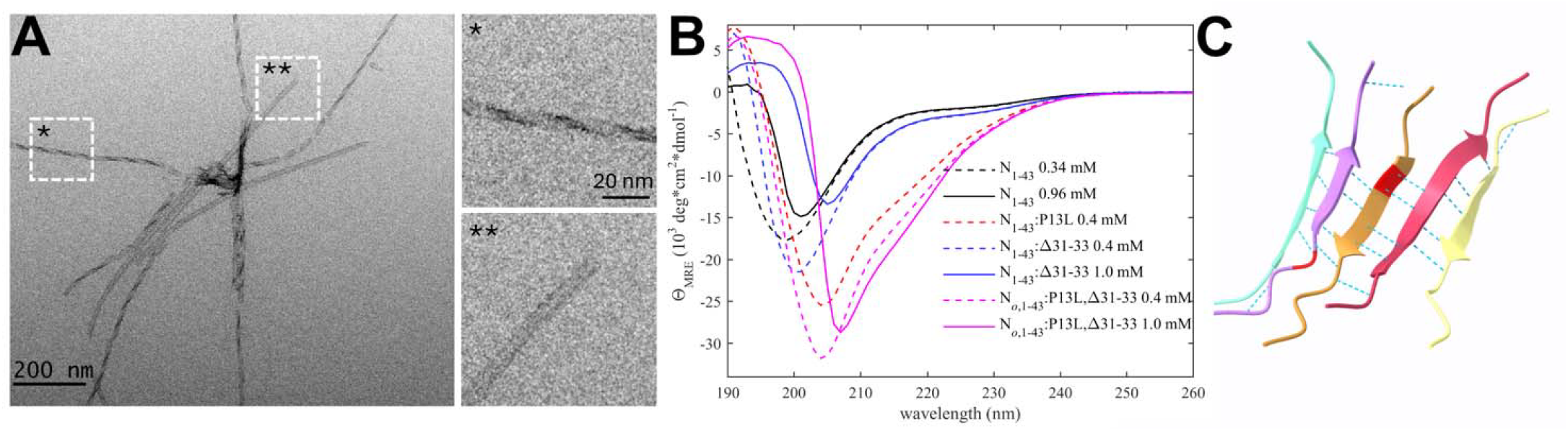
The P13L mutation creates self-association interfaces in the N-arm through stabilization of β-sheets. **(A)** Electron micrograph of negatively stained omicron N-arm N_ο_,_1-43_:P13L,Δ31-33 after equilibration at 10 μM in 20 mM HEPES, 150 mM NaCl, pH 7.50. The magnified regions are examples of twisted (*) and straight (**) fibrils. **(B)** CD spectra of N_1-43_:P13L (red), N_1-43_:Δ31-33 (blue), and Omicron N_ο_,_1-43_:P13L,Δ31-33 (magenta) at 0.4 mM (dashed) and 1.0 mM (solid), in comparison with ancestral N-arm (black). **(C)** Subset of ColabFold prediction of multimers of N_10-20_:P13L highlighting hydrogen bonds. The P13L residue is highlighted in red in the middle peptide.

Based on NMR studies of the N-arm, Zachrdla *et al*. previously reported a propensity for residues 13-19 to transiently populate extended/β-structure-like conformations (Zachrdla et al., 2022), which we hypothesized may be strengthened through the P13L mutation. To test the formation of β-sheet structure in N:P13L we carried out CD experiments of ancestral N-arm N_1-43_, and mutants carrying the P13L mutation, or Δ31-33, or both in the Omicron variant N_1-43_:P13L,Δ31-33 (**Figure 2B**). While the ancestral N-arm at ≈1 mM (≈4.6 mg/ml) concentrations exhibits CD spectra with a minimum at ≈200 nm typical of disordered conformations (black), the Omicron N-arm has a significantly higher structure content (magenta), consistent with β-sheets, as revealed by the strong negative mean residue ellipticity in the 210-220 nm range. Interestingly, diluting the 1 mM sample (solid) to a concentration of 0.4 mM (dashed) reveals a large shift in the far-UV spectra from positive to negative ellipticity at ≈200 nm, as well as a shift in the minimum to lower wavelengths, both indicative of a significant increase of disorder upon dilution. This is consistent with the stabilization of β-sheets in a reversible, strongly cooperative self-association process with an effective K_D_ in the high μM to low mM range. Dissecting the origin of the increased β-sheet content, it is apparent that the majority of the effect arises from the P13L mutation (red) alone, with a minor contribution by Δ31-33.

Finally, confirming the interpretation of the EM images and the CD data, as well as the β-structure propensity reported from NMR data (Zachrdla et al., 2022), the structural prediction of N_10-20_:P13L in ColabFold displayed oligomers with stacking β-sheets of residues 12-18 with typical hydrogen bond patterns (**Figure 2C**), whereas ancestral peptides did not lead to well-organized structures (**Supplementary Figure S4**). Analogous predictions of N-arms with two other frequent mutations, P13S and P13T, did not lead to β-sheet structures as in P13L.

While this self-association interface in the P13L N-arm is weak and its direct observation in biophysical experiments requires mM concentrations, which far exceed average intracellular concentration of N, such weak interactions can become highly relevant physiologically when high local concentrations are prevailing, for example, when the disordered extension is preconcentrated while tethered within macromolecular assemblies as in the RNP, or in macromolecular condensates.

### Enhanced oligomerization of the leucine-rich sequence through cysteine mutations

The protein-protein interaction interfaces driving LRS self-association have been studied in great detail in biophysical experiments and MD simulations for the ancestral molecule and several mutations in the mutational landscape (Zhao et al., 2022). A key feature is a pattern of hydrophobic residues that can combine to form a transient helix creating a hydrophobic surface stretching from ≈222 to 234 on one side of the helix. We have previously discovered the self-association of the LRS after analysis of the mutational landscape (Zhao et al., 2022) and reports of a conspicuous defining mutation N:G215C in Delta variants that correlated with its rise in 2021 among clades with identical spike mutations (Marchitelli et al., 2021; Stern et al., 2021; Zhao et al., 2022) . As deduced from MD simulations, the cysteine at position 215 is located at the base (N-terminal end) of the transient helix and, through its lower flexibility than the glycine, serves to redirect the adjacent upstream disordered residues such that helices are more prone to form stabilizing coiled-coil interactions (Zhao et al., 2023, 2022). As shown experimentally by sedimentation velocity analytical ultracentrifugation (SV-AUC) in reducing conditions, this enhances non-covalent self-association of LRS peptides as well as full-length N dimers by 2-3 orders of magnitude (Zhao et al., 2023, 2022). Covalent disulfide bonds in the LRS in non-reducing conditions were found to further promote LRS oligomerization. However, there is no conclusive data yet whether covalent bonds in the LRS occur *in vivo*, or any G215C effect is entirely non-covalent due to the significant strengthening of LRS helix oligomerization (see **Discussion**). In any event, the G215C mutation leads to enhanced assembly in a VLP assay (Zhao et al., 2024), and as shown in reverse genetics experiments, *in vivo* confers a significant replication advantage and an altered virion morphology (Kubinski et al., 2024).

Here we studied a mutation of G214, which in the mutational landscape exhibits a similar mutation pattern as G215. In particular, we focus on an independent introduction of a cysteine in the LRS that occurred in the Lambda variant, prevalent in South America in 2020-2021 (Wink et al., 2022), with the defining N-protein mutations P13L, R203K/G204R, and G214C (**Table 1**). (We will adopt a nomenclature where the complete set of defining mutations of a variant will be referred to by its Greek letter, i.e., N:P13L/R203K/G204R/G214C is N_λ_, and analogously the set of Omicron mutations N:P13L/Δ31-33/R203K/G204R are referred to as N_ο_; see **Table 1**). The effect of the G214C mutation is unknown. Due to the close proximity of 214 and 215, we asked whether it enhances LRS self-association similarly to G215C, and to this end first synthetized a LRS peptide comprising N_210-246_:G214C. Unexpectedly, unlike the chemically identical N_210-246_:G215C peptide, it exhibited low solubility. This prohibited its characterization at sufficiently high concentrations for study of self-association by SV-AUC, which required in excess of ≈0.4 mM for N_210-246_:G215C peptides to display oligomers. However, dynamic light scattering (DLS) revealed the presence of N_210-246_:G214C complexes with hydrodynamic radii ranging from 6 to 40 nm (in comparison to 1-2 nm for N_210-246_:G215C (Zhao et al., 2022)) in reducing conditions, and slightly larger in non-reducing conditions (**Supplementary Figure S5**). For N_210-246_:G214C a cumulant analysis results in radii of 8.8 nm and 10.6 nm and polydispersity indices of 0.40 and 0.35 for reducing and non-reducing conditions, respectively. This shows that while G214C also strengthens the LRS self-association interface, it exhibits different properties compared to G215C.

To gain more insight in the different behavior of the cysteine LRS mutants we carried out MD simulations of monomeric and trimeric LRS peptides N_210-246_. As shown in **Figure 3**, both cysteine mutants extend the helix by stabilizing the flexible GG motif into a well-defined α-helix turn. Importantly, however, the sulfhydryl groups in position 214 *vs 215 assume* different orientations relative to the hydrophobic patch serving as the oligomerization interface. Under reducing conditions, this has the potential to also cause variation in physicochemical properties of resulting oligomers, e.g., through repositioning of the adjacent glutamate E216 and resulting changes in the surface electrostatic potential, and to thereby alter the oligomerization scheme (**Supplementary Figure S6**). It can be expected that under oxidative conditions the differences between 214C and 215C are exacerbated due to their different relative orientation of the hydrophobic interface of the helix and the sulfhydryl groups, leading to different oligomeric states and different phase separation properties.

**Figure 3.**
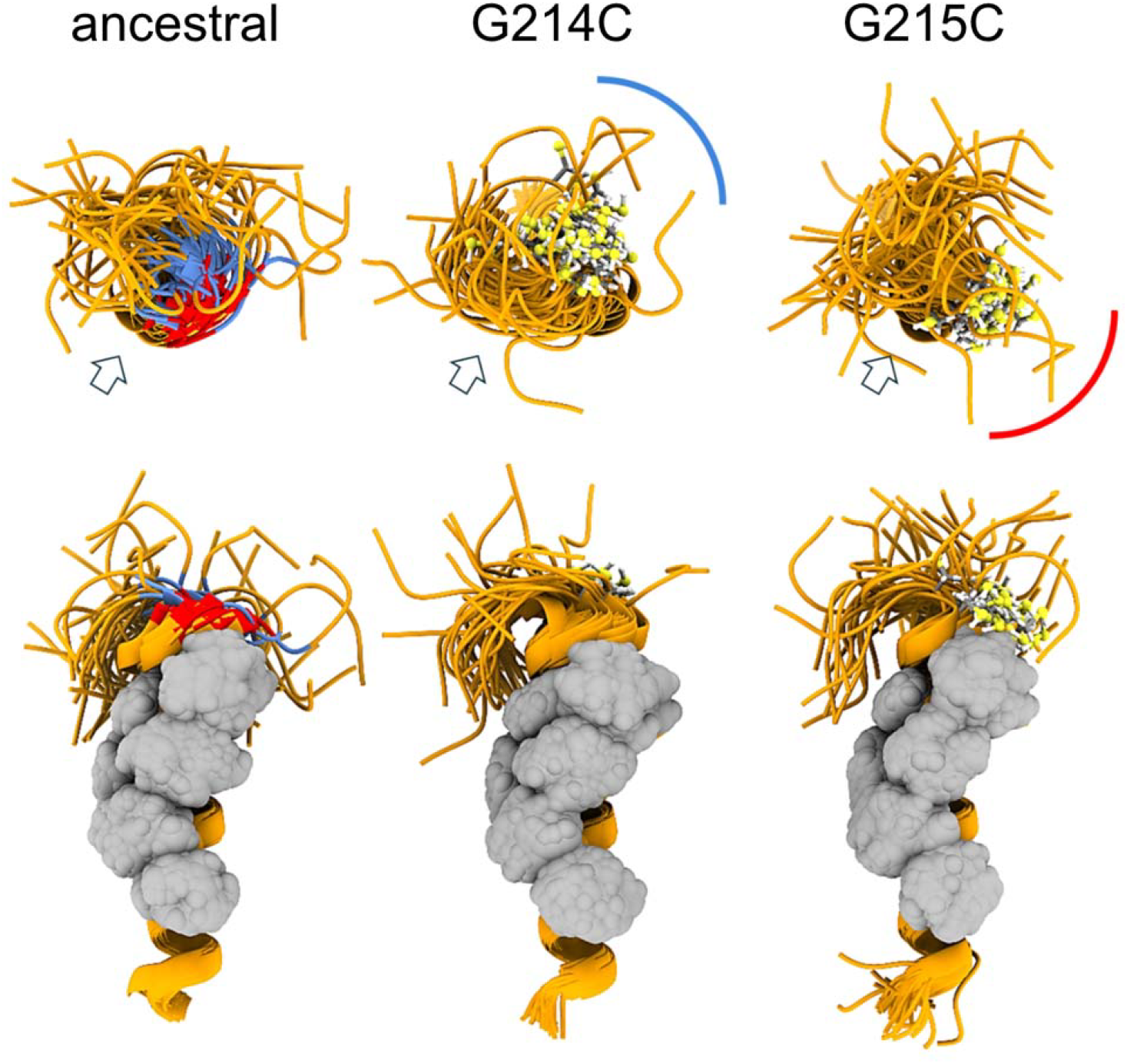
MD simulation of ancestral LRS and comparison with 214C and 215C mutants of Delta and Lambda variants. Snapshots at equal time intervals (4 ns) taken from the 200-ns MD simulations of the ancestral peptide N_210-246_ and single-point G→C mutants (all monomers were oriented by overlying the helical region and displayed in orthosteric view). The upper row shows a view from the N-terminus side (with the helix axes perpendicular to the plane of the figure), while the lower row presents a side view (with axes on the plane). For clarity, the disordered C-terminal segment (residues 236-246) has been removed. The glycine and cysteine residues at positions 214 and 215 (colored blue and red, respectively, in the ancestral peptide and rendered as ball-and-stick in the mutants) restructure the flexible backbone around the GG motif into a well-defined α-helix turn, directing the sulfhydryl group in specific orientations (illustrated schematically by the blue and red arches). These orientations can be quantified relative to the Leu-rich central region (indicated by arrows and rendered as gray van der Waals spheres), which forms the hydrophobic interfaces of the oligomers (Zhao et al., 2023). Under reducing conditions, this reorientation of the N-terminus relative to the helix can influence helix binding during the early stages of oligomerization or alter the conformation and physicochemical properties of the resulting oligomers, as illustrated in **Supplementary Figure S6**.

### Impact of enhanced self-association interfaces on the oligomerization of full-length N

In principle, since N-protein is a constitutive dimer coupled in the CTD, binding interfaces in the LRS and N-arm may form intra-dimer bridges simply further stabilizing the dimeric state. Indeed, coarse-grained molecular simulations of N-protein dimers have revealed large conformational fluctuations where the LRS, for example, can make rare intra-dimer contacts (Różycki and Boura, 2022). On the other hand, given the large flexibility of the disordered chain, the same interfaces may lead to higher oligomers if they establish one or two inter-dimer bridges across different dimers, generating different classes of tetramers. We would expect a concentration-dependent probability of such inter-dimer contacts for N-protein in solution.

Experimentally, in the absence of nucleic acid ligands, ancestral N-protein at low μM concentrations is essentially dimeric with only hints of reversible higher-order oligomers (Yu et al., 2005; Zhao et al., 2021). With the strengthened LRS helix stability through the G215C mutation, in reducing conditions, we observe reversible tetramerization with a K_D_ of 1.0 (0.7 – 1.5) μM (**Figure 4A**), consistent with previous work (Zhao et al., 2022). We made analogous observations by SV-AUC for N_λ_ (containing the G214C mutation) in reducing conditions, demonstrating it similarly can form reversible inter-dimer bonds (**Figure 4A**). Clearly, strengthened LRS helices can alter the N-protein oligomeric state by crosslinking dimers into tetramers.

**Figure 4.**
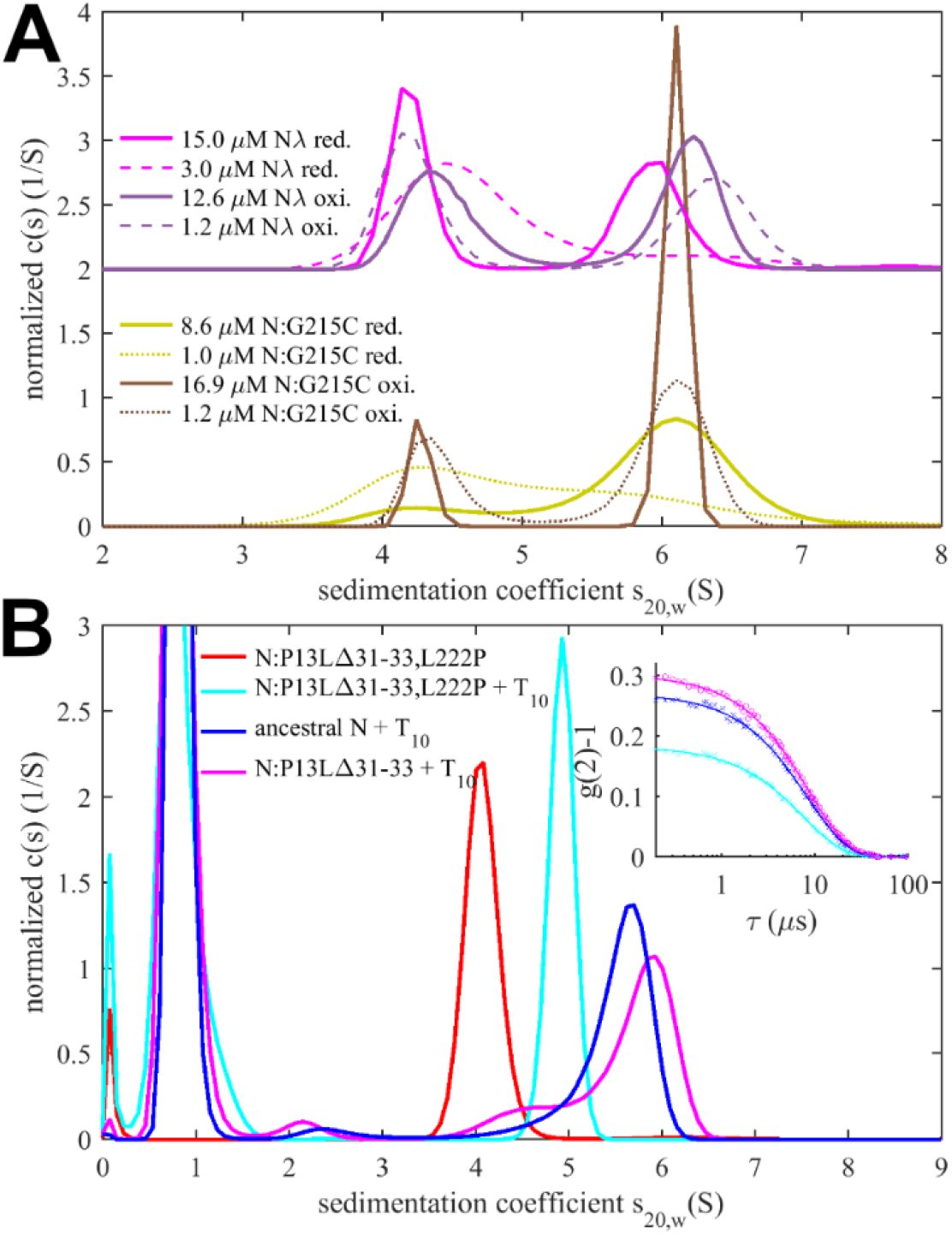
Impact of mutations in self-association interfaces on oligomeric state. **(A)** Sedimentation coefficient distributions of cysteine mutants N:G215C (reduced in yellow) and N:G215C* (oxidized in brown), as well as reduced N_λ_ (reduced in magenta) and N_λ_* (oxidized in violet), with N_λ_ data offset by 2. For each sample data were acquired at high (solid lines) and low (dashed lines) concentration. **(B)** Sedimentation coefficient distributions of 2 μM N:P13L,Δ31-33, ancestral N, and N:P13L,Δ31-33,L222P in the presence of 10 μM T_10_ in low salt buffer. The inset shows DLS autocorrelation data of the same samples (symbols) and single-species fits (lines).

To study the potential impact of disulfide bonds on N-protein oligomeric state, we prepared oxidized full-length N:G215C and N_λ_ by extensively dialyzing the reduced protein in TCEP-free buffer while purging air through the dialysate for gentle passive oxygenation. This oxidized sample is referred to as N:G215C* and N_λ_* . Using a DTNB assay we assessed the percentage of free sulfhydryl groups to be ≈30% for both N:G215C* and N_λ_*, respectively. Since the LRS mutation provides the sole cysteine in N-protein, we concluded that the majority of N:G215C* and N_λ_* is disulfide-linked in the LRS, which was confirmed by non-reducing SDS-PAGE (**Supplementary Figure S7**). Due to the non-covalent dimerization in the CTD as well as non-covalent tetramerization *via* LRS interfaces (see above), these samples may be complex mixtures of dimers and/or higher oligomers with different patterns of covalent and non-covalent intra-and inter-dimer LRS interactions. Using SV-AUC we established that close to half of both the N:G215C* and N_λ_*sample is tetrameric at low μM concentrations (**Figure 4A**). Because oxidation significantly increases the tetramer population, we conclude that one or two covalent inter-dimer bonds can form that enhance N-protein oligomerization.

In contrast to the modulation of the coiled-coil LRS interfaces, the *de novo* creation of the N-arm self-association interface through beta-sheet interactions enabled by P13L cannot be readily observed in full-length N-protein at low μM concentrations. Similar to the ancestral LRS interface, it provides only ultra-weak binding energies that require mM concentrations to significantly populate oligomers. This is fully consistent with the previous observation by SV-AUC that neither N:P13L,Δ31-33 nor N_ο_ with the full set of Omicron mutations show any significant higher-order self-association at low μM concentrations, whereas at high local concentrations – as observed in phase-separated droplets – they can modulate and cooperatively enhance self-association processes (Nguyen et al., 2024). (If fact, P13L can substitute for the LRS promoting LLPS, as observed in the rescue of LLPS by N:P13L,Δ31-33/L222P mutants whereas N:L222P LRS-abrogating mutants are deficient in LLPS.) Another process that increases the local concentration of N-arm chains is the tetramerization of full-length N-protein. As described earlier, occupancy of the NA-binding site in the NTD allosterically promotes self-assembly of the LRS into higher oligomers (Zhao et al., 2021). We hypothesized that these oligomers may be cooperatively stabilized by additional N-arm interactions in P13L mutants.

To this end, we carried out SV-AUC and DLS experiments of 2 μM ancestral N, and the full-length N-arm mutant N:P13L,Δ31-33 in the presence of a short oligonucleotide T_10_ that occupies the NTD binding site but is too short for scaffolding multiple N-protein dimers (Zhao et al., 2021) (**Figure 4B**). Low salt buffer (10 mM NaCl, 2.7 mM KCl, 10.1 mM Na_2_PO_4_, 1.8 mM KH_2_PO_4_, pH 7.4) in this experiment ensures maximal occupancy of the NTD binding site for nucleic acid by 10 μM T_10_, which under these conditions has a K_D_ below 0.1 μM (Zhao et al., 2021). In a control experiment by SV-AUC we measured the affinity of the T_10_ oligonucleotide for ancestral N and N:P13L,Δ31-33 and observed no significant difference in moderate ionic strength (**Supplementary Figure S8**). Ancestral N protein in the absence of oligonucleotide sediments at ≈4.0 S, reflecting its dimeric state afforded by the CTD dimerization domain (Zhao et al., 2021). When LRS oligomerization is abrogated through introduction of a L222P mutation, binding of T_10_ to the N-protein dimer still induces a conformational change and increases its *s*-value to ≈4.9 S (Zhao et al., 2023). With the native LRS in the ancestral N this T_10_ -ligated state allows LRS oligomerization, which can be observed through the formation of a reaction boundary in SV-AUC with an *s*-value of ≈5.6 S, reflective of a mixture of dimers and tetramers in rapid exchange relative to the time-scale of sedimentation (Schuck, 2010; Zhao et al., 2023). As a control, we reproduced this previously reported result (**Figure 4B**, blue). Introduction of the N-arm mutations N:P13L,Δ31-33 causes a further increase of the *s*-value of the reaction boundary to ≈5.8 S, indicating an increase in the tetramer stability (**Figure 4B**, magenta). This stabilization of the tetramer is corroborated independently by an increase in the average hydrodynamic radius of N:P13L,Δ31-33 in mixture with T_10_ (5.51 nm) relative to ancestral N (5.37 nm) or the LRS mutant N:P13L,Δ31-33,L222P (4.84 nm) (**Figure 4B inset**). Thus, the N-arm mutation clearly strengthens inter-dimer interactions, even though the added binding energy is too weak to produce detectable tetramer populations at micromolar concentrations by itself. In principle, allosteric interactions between the distant disordered N-arm and the LRS in the disordered linker might exist that cause enhanced tetramerization of N:P13L,Δ31-33 with occupied NA-site in the NTD. However, a more parsimonious explanation is that the additional self-association interface in the N-arm created by P13L makes inter-dimer contacts that add to the separate oligomerization of the LRS helices in stabilizing tetramers.

### Mutation effects on RNP assembly and stability

As the assembly of RNPs requires the concerted effect of several binding interfaces, we asked whether the enhanced LRS coiled-coil stability and the novel N-arm self-association interface impact the RNP stability. To examine this experimentally we carried out *in vitro* RNP assembly experiments using the assay developed previously by the Morgan laboratory (Carlson et al., 2022, 2020). As mentioned above, it is based on the observation that mixtures of N-protein with stem-loop RNA from the viral 5’-UTR at low μM concentrations readily form polymorphic ribonucleoprotein complexes that match in size, symmetry, and RNA content what would be expected for the assembled RNPs in virions (Carlson et al., 2022, 2020). The use of stem-loop SL7 as RNA substrate helps to minimize structural polydispersity arising from variable secondary structure elements. We embarked on the experimental roadmap introduced previously (Zhao et al., 2024) consisting of SV-AUC experiments that hydrodynamically resolve RNPs in dynamic assembly equilibrium in solution as fast-sedimenting reaction boundaries (Schuck, 2010; Schuck and Zhao, 2017), in combination with complementary mass photometry (MP) experiments that can resolve populations of different protein/RNA complexes and their dissociation products at sub-μM concentrations through interferometric sizing of single-molecule surface adsorption events (Wu and Piszczek, 2021).

Sedimentation coefficient distributions of ancestral and different mutant full-length N at 3 μM in the presence of 3.4 μM SL7 are shown in **Figure 5A**. For reference, under these conditions the RNPs of the ancestral N-protein form a reaction boundary with a weighted average sedimentation velocity of 19.7 S (black). Under otherwise identical conditions, significantly faster sedimentation can be discerned for N:P13L,Δ31-33 (20.2 S, red). Faster sedimentation can reflect an increase in size of the complexes, and/or increased populations and lifetimes of complexes in dynamic equilibrium, i.e., higher affinity and stability. By contrast, N:R203K/G204R exhibits a slightly lower *s*-value of 19.5 S (blue). Both R203K/G204R and P13L,Δ31-33 combine in the set of defining Omicron mutations, N_ο_, which produces RNP boundaries at 20.5 S (cyan). It appears the N-arm mutations can more than compensate for the loss of RNP stability through the R203K/G204R mutation in the linker. The introduction of the LRS cysteine in N:G215C is nearly as effective, with an *s*-value of 20.4 S (orange). Finally, the largest increase in sedimentation velocity can be discerned for the combination of P13L/R203K/G204R with the LRS enhancing cysteine G214C in N_λ_ with 21.0 S (magenta) (see **Table 1**).

**Figure 5.**
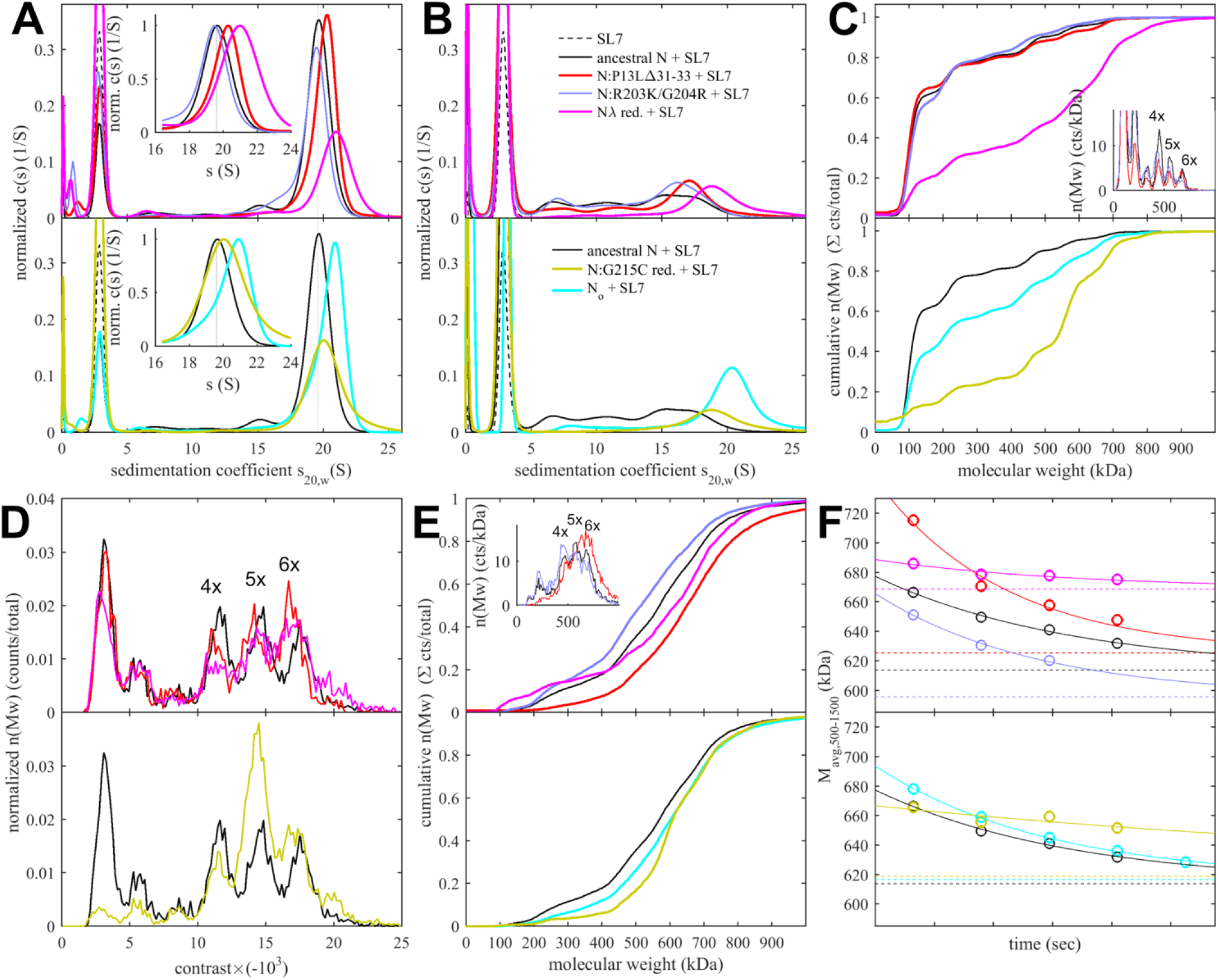
Size distributions and stability of ancestral and mutant RNPs. Show are SV (A,B) and MP data (C-F) for mixtures of N-protein with stem-loop RNA SL7 in molar ratio of 1(N):1.15(SL7) at high concentration of 3 μM (A,D) or low concentration of 0.3 μM (B,C,E,F) in reducing buffer conditions. All panels use the same color scheme for N-protein: ancestral (black), N:P13L,Δ31-33 (red), N:G215C (orange), N_ο_ (cyan), N:R203K/G204R (blue), N_λ_ (magenta). All panels are subdivided in two plots for clarity, with each showing ancestral trace in black for comparison. **(A)** Sedimentation coefficient distributions of mixtures equilibrated at high concentration, with reaction boundary peaks magnified in the inset. For reference, the ancestral RNP s-value is drawn as dotted vertical line. Absorbance data are recorded at 260 nm and are weighted by SL7 content of sedimenting species. Higher reaction boundary s-values signify greater affinity or lifetime of the mutant RNPs. **(B)** Sedimentation coefficient distributions of the same samples as in (A), tenfold diluted and equilibrated, highlight dissociation of most RNPs into a range of intermediate size complexes. **(C)** MP experiments of equilibrated 0.3 μM mixtures. The measured number distributions are presented as cumulative distributions, which display higher percentages of large species as shifts to the right. Most samples are largely dissociated into dimers, with remaining peaks corresponding to populations of dimer to hexamers of N_2_/SL7_2_ subunits (as highlighted in the differential distributions in the inset). As an example for the resolution of distinct species, the inset shows the differential distribution (histogram) for N:R203K/G204R (blue), ancestral N (black), and N:P13L,Δ31-33 (red), with the peak labels indicating the number of N-dimer/2SL7 subunits. **(D)** Mass distributions acquired in stopped-flow configuration applied to 3 μM mixtures. Larger (negative) contrasts correspond to higher molecular weights, with major peaks corresponding to species containing 1, 4, 5, and 6 N_2_/SL7_2_ subunits. (E) For kinetic experiments, mass distributions were acquired in different time intervals after tenfold dilution of 3 μM mixtures, here showing data collected from 3 sec to 23 sec. **(F)** Number-average molecular weights of assembled RNPs between 500 and 1500 kDa observed in consecutive 20 sec data acquisition intervals after tenfold dilution of 3 μM mixtures (circles). The dashed horizontal lines are number-averages determined from the equilibrated 0.3 μM mixtures in **(E).** The solid lines are a best-fit single exponentials constrained to decay to the measured equilibrium values, yielding RNP lifetimes listed in **Table 1**.

When these samples are diluted and equilibrated at tenfold lower concentration, the RNPs are largely dissociated, as may be discerned from the reduced amplitude of the rapidly sedimenting peak and significant population of species with *s*-values between 7 and 18 S (**Figure 5B**). This highlights the cooperative oligomerization of the N-dimer/2SL7 subunits observed previously (Zhao et al., 2024) . Interestingly, the only partial dissociation of N_ο_, N:G215C, and N_λ_ RNPs suggests the existence of a subpopulation with higher stability. By contrast, the sedimentation coefficient distribution of RNPs from 0.3 μM N:P13L,Δ31-33 is more similar to that of the ancestral N-protein and N:R203K/G204R, suggesting the augmented RNP population caused by combining these mutations is kinetically not as stable and the contributions from N-arm interfaces are more transient.

Side-by-side to the SV experiments, MP measurements were carried out on the same samples to shed more light on the mass distribution of the RNP particles. For discrimination of individual surface adsorption events on the coverslip, the sample concentrations cannot exceed 0.3 μM protein with 0.34 μM SL7. As shown in **Figure 5C**, this results in a ladder of oligomers of N-dimer/2SL7 subunits ranging from ≈120 kDa (a single N-dimer/2SL7 subunit) to ≈700 kDa (a hexamer of subunits) (Carlson et al., 2022; Zhao et al., 2024). While this is analogous to the solution data in **Figure 5B**, for the detailed comparison of SV and MP distributions their different signal weights should be considered: where MP counts individual particles producing a number distribution, SV records a signal proportional to mass, and therefore is strongly skewed toward larger particles compared to MP. Also, in a trade-off between resolution and susceptibility to systematic errors, there is a potential for chemical properties to bias the surface adsorption, and for some adventitious experimental variation originating from the glass substrates. Nonetheless, we obtained results qualitatively consistent with the SV-AUC data, where N:P13L is largely dissociated like ancestral N-protein and N:R203K/G204R at 0.3 μM; and at the same time, N_ο_, N:G215C, and N_λ_ RNPs retain more of the oligomeric state. Interestingly, N:G215C shows the least fully dissociated dimer and retains more of a pentameric species, which is in contrast to the other cysteine-containing mutant N_λ_ (**Figure 5C**).

The ladder of oligomeric subunits resolved in MP poses the question whether the largely assembled state at 3 μM is uniform. Under these conditions SV-AUC shows a single RNP reaction boundary, but it is relatively broad, which would be equally consistent with a reaction boundary from a single complex in rapid association/dissociation exchange on the time-scale of sedimentation (< 1000 sec) (Schuck and Zhao, 2017), as with a polydisperse mixture of several unresolved large oligomers. In order to gain more insight in the mass distribution under these 3 μM conditions, for some constructs we employed a microfluidic accessory device for the MP instrument allowing rapid dilution with a dead time of < 0.1 sec to minimize RNP dissociation. The resulting data in **Figure 5D** again exhibit a ladder of oligomeric peaks, where the majority of N-protein assembled in a heterogeneous mixture of RNPs between tetramer and hexamer of subunits. Thus, the dissociation products after dilution in **Figure 5C** do not seem to originate from a single assembled complex. Furthermore, we observe characteristic differences in the oligomeric distribution between different constructs. Similar to the equilibrated lower concentration conditions, relative to ancestral N, the hexamer population is augmented for N:P13L,Δ31-33, N:G215C, and N_λ_. Notably there is again a prominent pentamer population for N:G215C.

Since the disassembly of RNPs after viral entry is another critical step in the viral life cycle, we aimed to probe the kinetic stability of the RNP complexes. To this end, we applied a modified pipettor-based sample application protocol where rapid dilution of the 3 μM mixture was followed by several consecutive 20 sec periods of data acquisition. The data from the acquisition immediately following the dilution (acquired between 3 – 23 sec) is shown in **Figure 5E**. It provides mass distributions showing substantial assembly into heterogeneous populations of RNPs. Qualitatively consistent with the 3 μM mixtures in SV-AUC, the N:R203K/G204R has a clear destabilizing effect on the RNP, whereas all mutants with enhanced binding interfaces exhibit higher populations of larger RNPs, with the greatest enhancement for N:P13L,Δ31-33. In order to focus on the kinetics of RNP dissociation we calculated the number average molecular weight of RNPs, which is plotted as a function of decay time in **Figure 5F** and empirically fitted as a single exponential decay attaining the separately measured equilibrium value measured at 0.3 μM (for distributions see **Figure 5C**, for number averages **Table 1**). While N:P13L,Δ31-33 produces the largest increase in RNP molecular weight between 0.3 μM and 3 μM, these RNPs have the shortest lifetime (τ = 44 sec *vs* 66 sec for the ancestral N), suggesting the added H-bond interactions in the N-arm to be rapidly reversible. On the other hand, RNPs of N:G215C have a lower average mass (being dominated by a pentameric oligomer of subunits), but show an increased kinetic stability (τ = 231 sec). RNPs of N_λ_ carrying both an LRS cysteine and the N-arm mutation are simultaneously of higher average molecular weight and persist most upon dilution (with a best-fit RNP equilibrium level of 669 kDa). (**Figure 5F**).

Even though it is still unclear whether disulfide bonds of N cysteine mutants form *in vivo*, we were curious about the impact of disulfide-linked oligomers of the cysteine mutants on their RNP structure and stability in our biophysical assembly model, and carried out analogous experiments with the substantially oxidized protein preparations depicted above in **Figure 4A**. As shown in **Figure 6A**, sedimentation coefficient distributions of 3 μM oxidized N:G215C* and oxidized N_λ_* (carrying 214C) in assembly mixtures with SL7 show faster reaction boundaries when oxidized as compared to their reduced form, with 20.8 S *vs 20*.*4 S for* N:G215C, and 21.5 S *vs 21*.*0 S for* N_λ_. Upon dilution the RNPs of oxidized N:G215C* protein dissociate similar as those of reduced N:G215C. For oxidized N_λ_ RNPs dissociation is less, but still significant. The molecular weight distributions in MP measured in the first 20 sec after rapid dilution of the 3 μM stock (**Figure 6B**) show largely assembled mixtures of tetramers, pentamers, and hexamers of the N-dimer/2SL7 subunits, with a large increase in the population of hexameric RNPs in oxidized *vs reduced for*ms. The tail of even higher molecular weight species is enhanced, pointing to subpopulations of heptameric species especially for oxidized N_λ_ RNPs. Interestingly, the preference of pentameric RNPs formed by reduced N:G215C is absent after oxidation. N_λ_* exhibits the largest population of hexamers observed in any sample, in agreement with the highest *s-value of t*he reaction boundary in SV. As indicated by the time-course of RNP dissociation after tenfold dilution (**Figure 6C**), oxidation enhances the average molecular weight of the RNPs but not their lifetime. We conclude that disulfide-linked N-protein tetramers can incorporate into and aid in the formation of RNPs, modulating their preferred oligomeric state, and that cumulative effect of the N-arm mutation P13L and the LRS cysteine persists for disulfide-linked protein.

**Figure 6.**
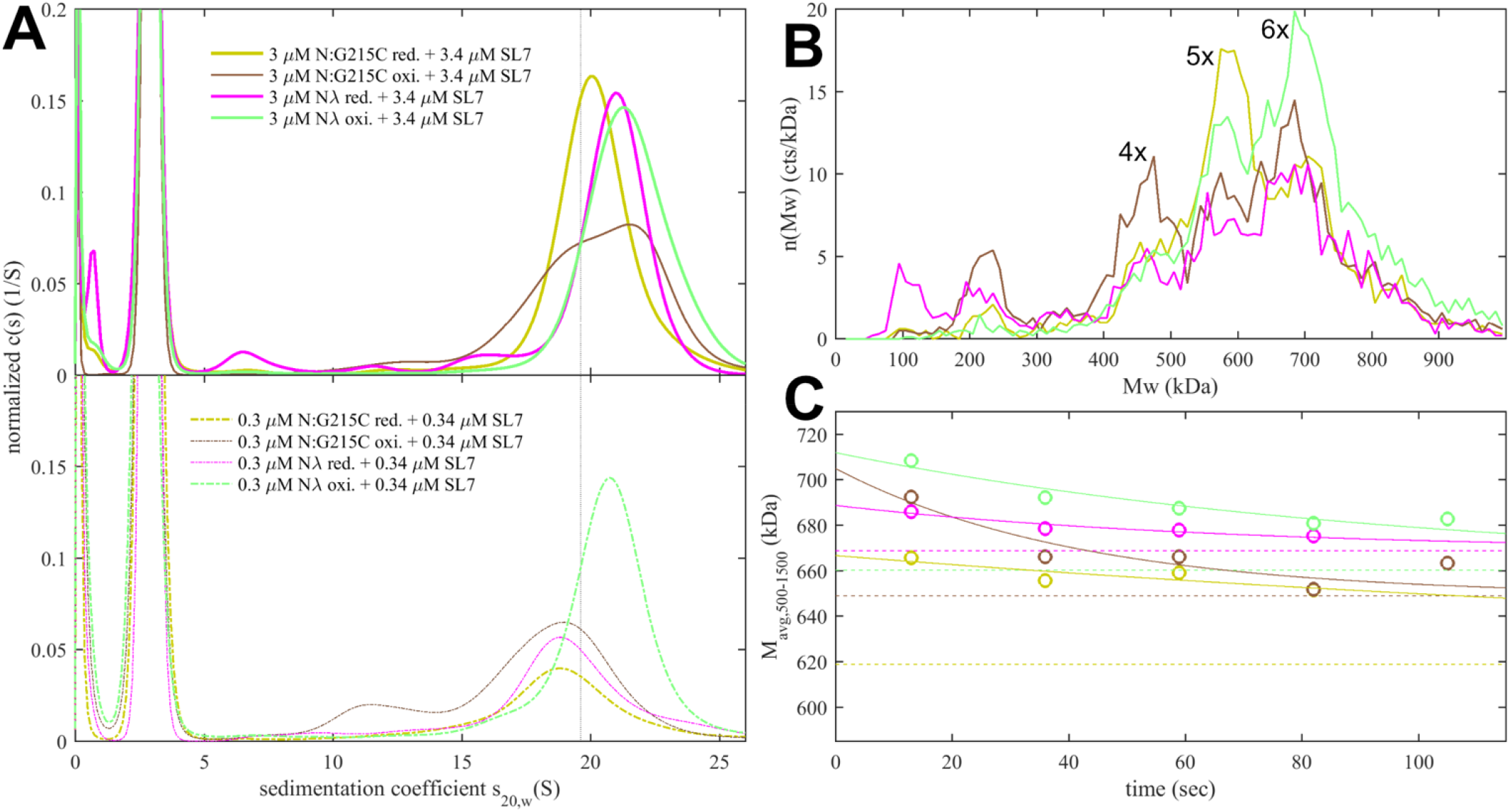
Impact of LRS disulfide bonds on the size and stability of RNPs. N-protein with cysteine mutations in the LRS were oxidized to form disulfide-linked oligomers (as shown in **Figure 4A**) and mixed with stem-loop RNA SL7 in molar ratio of 1(N):1.15(SL7). Shown are data for oxidized N:G215C* (brown) and oxidized N_λ_* (green), and for comparison, reduced N:G215C (yellow) and reduced N_λ_ (magenta). **(A)** Sedimentation coefficient distributions at 3 μM (upper panel) and 0.3 μM (lower panel) protein. **(B)** Molecular weight distributions in MP experiments of the same mixtures rapidly diluted to 0.3 μM protein, acquired from 3 – 23 sec after dilution, with peak labels reflecting the multiples of N dimer/2SL7 subunits. **(C)** Time-course of number-average RNP molecular weights between 500 and 1,500 kDa (circles), determined from rapid dilution experiments in (B) for consecutive 20 sec data acquisition intervals. The solid lines are best-fit single-exponential decays constrained to attain the separately measured equilibrium values at 0.3 μM protein (dashed lines), with lifetimes listed in **Table 1**.

#### Virus-like particle formation and infectivity

We asked whether the observed modulation of RNP size and stability by N mutations impacts the formation and infectivity of virus-like particles (VLPs) (Syed et al., 2021; Zhao et al., 2024). In this assay, producer 293T cells are co-transfected with plasmids for the four structural proteins S, E, M, and N of SARS-CoV-2, alongside a plasmid containing the viral packaging sequence T20 (nt 20080 – 22222, located near the 3’ end of ORF1ab) with a luciferase reporter gene, with a total length of 4,127 nt. This leads to assembly of VLPs, which are collected from the supernatant and applied to receiver cells transfected with entry factors ACE2 and TMPRSS2. Infected receiver cells then express the luciferase reporter and luminescence is measured, as an indicator for the combined efficiency of protein expression, VLP assembly in the producer cells, and entry into the receiver cells (Syed et al., 2021).

First, a control experiment was carried out with the N:L222P mutant previously shown to abrogate LRS oligomerization and RNP formation. Similar to a second control without N-protein plasmid, N:L222P produced very little luminescence relative to ancestral N (**Figure 7**). A positive control was N:R203M (green), which was previously shown to significantly enhance the signal of the VLP assay (Syed et al., 2021), and did so in the present VLP experiments. Similarly consistent with previous reports (Syed et al., 2021; Wu et al., 2021), R203K/G204R (blue) led to increased VLP signals. Of the mutations related to N-protein binding interfaces examined in the present work, neither P13L (red), nor Δ31-33 (light red), nor G215C (yellow) alone led to significant enhancement; only G214C (orange) produced a small but statistically significant enhancement. However, combination of the latter with P13L in P13L/G214C (brown) increased the gain, and further incorporation of the R203K/G204R double mutation to produce the full set of N_λ_ mutations (purple) substantially increased the measured luminescence. A similar observation was made for the combination of P13L/Δ31-33/R203K/G204R constituting the full set of N_o_ mutations (magenta). As expanded on in the **Discussion**, the failure to observe enhancement by P13L alone may be related to limitations of the VLP assay in sensitivity, including the restriction to a single round of infection, and protein expression levels.

**Figure 7.**
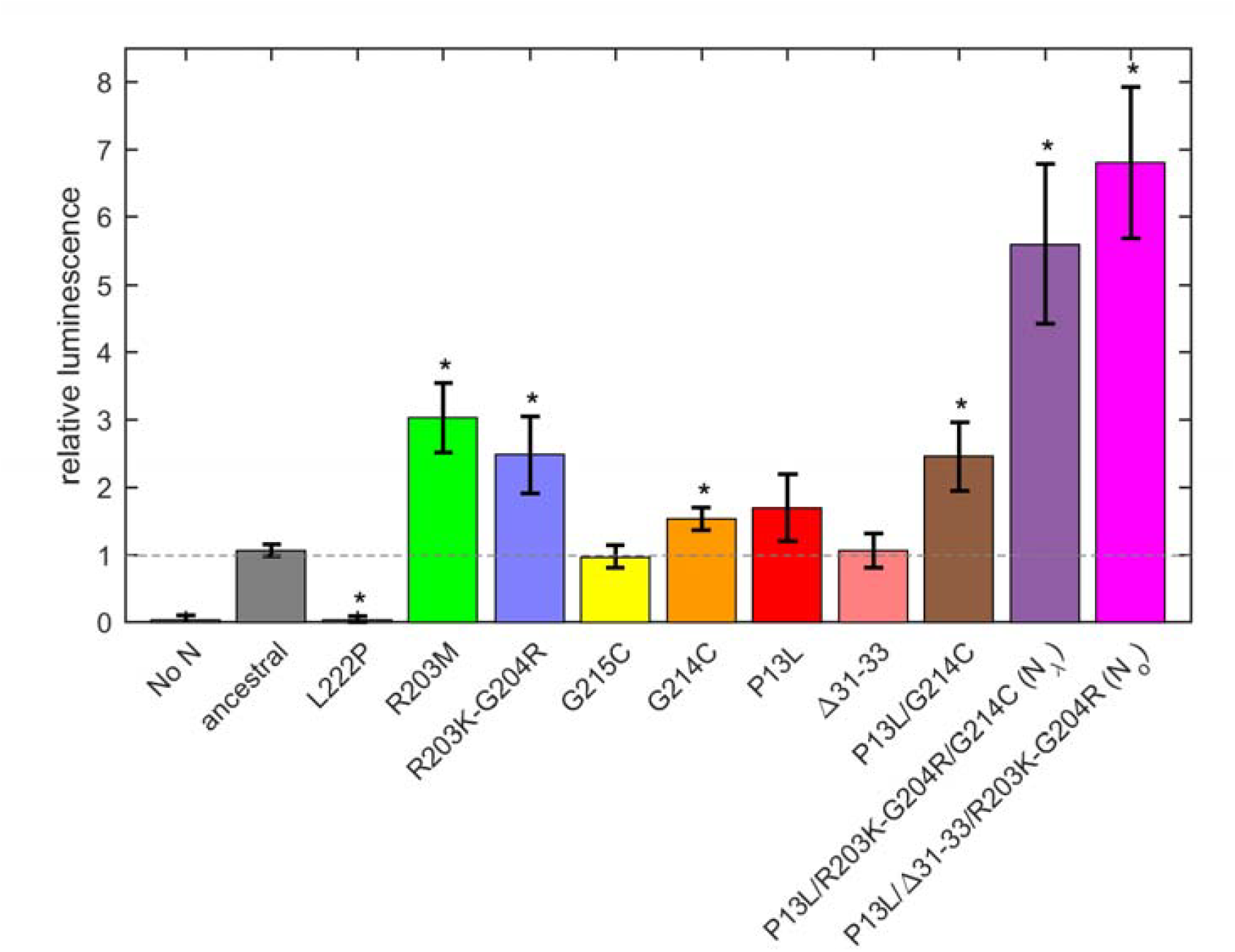
Mutation effect on packaging and cell entry in a VLP assay. Error bars are standard deviations from n = 4. Stars indicate significance (P > 0.95) of a two-sided Kolmogorov-Smirnov test comparing the control ancestral measurements with mutants.

### P13L mutation in N-protein promotes SARS-CoV-2 replication in cell lines

Finally, we characterized the fitness of P13L mutation by introducing it into a recombinant SARS-CoV-2 reporter virus expressing an mCherry gene fused to N *via* a P2A linker (C. Ye et al., 2020). We infected Vero-TMPRSS2 and A549-ACE2 cells at an MOI of 0.01 with Wuhan-Hu-1 (ancestral) and mutant viruses, and measured mCherry fluorescence as well as viral release into the supernatant at indicated time points post-infection. This revealed that the P13L mutant exhibited a stronger mCherry signal throughout the course of infection (**Figure 8A,B**) and generated more progeny virus strain in both cell lines (**Figure 8C**). These findings are consistent with P13L providing a fitness advantage over ancestral virus.

**Figure 8.**
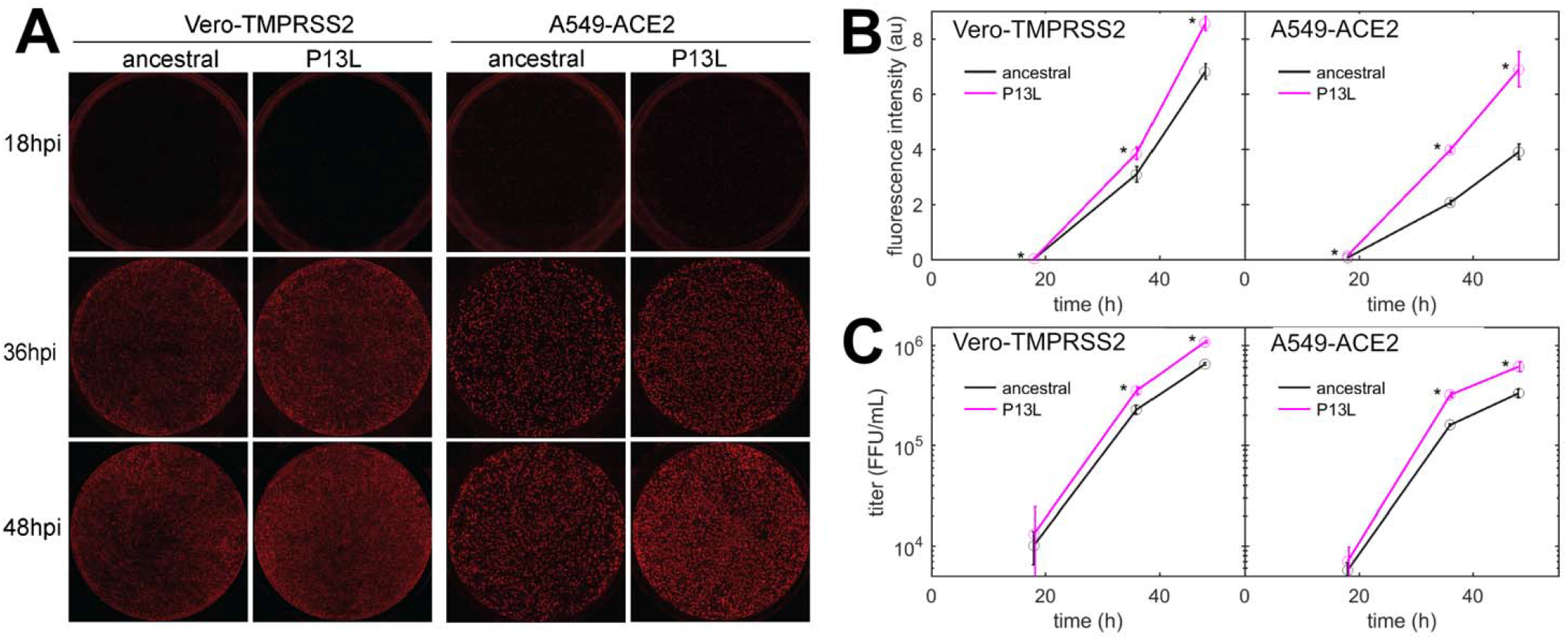
Replication kinetics of recombinant SARS-CoV-2 reporter viruses in cell lines. **(A)** Representative images of Vero-TMPRSS2 and A549-ACE2 cells infected with SARS-CoV-2 P13L or WT at different time points post-infection. **(B)** Quantification of fluorescence intensity from P13L and WT virus infections shown in (A). (C) Viral titers in the supernatant from infected cells. Error bars are standard deviations (n = 3), and stars indicate significant differences on a P=0.95 confidence level.

## Discussion

Rapid evolution of protein binding interfaces has frequently been observed in viral protein complexes, notably in the virus-host interface, including viral surface glycoproteins as well as ribonuclear proteins and non-structural proteins, with fitness advantages being accomplished, for example, through reshaping the binding interfaces, modulating protein structural dynamics, or altering physicochemical properties (Barozi et al., 2022; Evseev and Magor, 2021; Focosi et al., 2024; Planchais et al., 2022; Rochman et al., 2022). Also, entirely new interactions can arise through the viral mimicry of eukaryotic short linear motifs as a result of frequent mutations in the viral protein intrinsically disordered regions, which can greatly augment the virus-host interface (Davey et al., 2015; Schuck and Zhao, 2023). The mutations we have studied here are of a different category, impacting the interactions among viral proteins that enhance viral multi-protein complexes. Previous examples include the intra-host diversity of polymerase subunit interfaces in H5N1 influenza viruses (Welkers et al., 2019). While such mutations are not directly targeted towards the host, they may still contribute to host adaptation, or be balancing other mutation effects in an epistatic network, conceivably involving modulation of local effective protein concentrations (P. Li et al., 2023). Irrespective of their complete context, they can provide valuable insights in viral protein mechanisms.

Specifically, we have described three different mutations of SARS-CoV-2 N-protein that, in convergent evolution, strengthen the formation of RNPs and enhance viral assembly. N:G214C, N:G215C, and N:P13L have been independently introduced (as highlighted in the phylogenetic trees **Supplementary Figure S9-11** generated in Nextstrain (Hadfield et al., 2018)), and persisted in the defining set of mutations in their respective variants of concern (Lambda, Delta, and Omicron, respectively). We have shown here that N:P13L confers a fitness advantage in cell lines, and similarly, N:G215C was shown by Kubinski *et al*. (Kubinski et al., 2024) to impart improved viral growth. This correlates well with our results studying their molecular mechanisms.

For both of the cysteine mutants, molecular dynamics simulations and biophysical studies show how cysteines augment self-association interfaces by extending and redirecting the transiently formed helical coiled-coils in the intrinsically disordered LRS, which play a central role in the assembly of RNPs. By contrast, for N:P13L, unexpectedly, the evolution of RNP stability goes beyond modulation of a previously existing binding interface, and instead we observe the *de novo* formation of an additional dynamic self-association interface in the distant disordered N-arm through the stabilization and stacking of transient β-sheets, that we hypothesize cooperatively contributes to the stability of RNPs. Even though the solution affinity of the N-arm P13L interface is ultra-weak, the average local concentration of N-arm chains across the RNP volume (in a back-of-the-envelope calculation assuming a ≈14 nm cube (Klein et al., 2020) with a dodecameric N cluster) is ≈7.4 mM, such that disordered N-arm peptides could well create populations of N-arm clusters stabilizing RNPs through this interface.

However, besides the RNP-stabilizing mutants we have also observed unexpected RNP destabilization by the ubiquitous R203K/G204R double mutation, which may be caused by the introduction of additional charges close to the self-association interface in the LRS. In our experiments, this destabilization is more than compensated for by the P13L mutation. (Another scenario where ultra-weak interactions can have a critical impact is in molecular condensates. We previously reported the suppression of LLPS by the R203K/G204R mutation, which is rescued by the additional P13L/Δ31-33 mutation (Nguyen et al., 2024). This is consistent with compensatory weak stabilizing and destabilizing impacts of weak interactions on the RNP observed here.)

We arrive at a picture of SARS-CoV-2 RNPs that is far from structurally well defined, matching the concept of fuzzy complexes (Wu and Fuxreiter, 2016). On a molecular level, large portions of the SARS-CoV-2 N-protein (the N-arm, C-arm, and linker) are intrinsically disordered and highly flexible (Cubuk et al., 2021; Różycki and Boura, 2022), which persists in the presence of bound nucleic acid (Cubuk et al., 2024; Guseva et al., 2021; Schiavina et al., 2022) . It appears that conformational freedom is also retained to a significant degree in the RNPs. This flexibility could be advantageous for accommodating various RNA secondary structures (Carlson et al., 2022; Landeras-Bueno et al., 2025), and favorably balance the energetic cost of RNP disassembly that is required immediately after viral entry. Also, this serves to accommodate significant sequence variation(Davey et al., 2011; Duro et al., 2015; Schuck and Zhao, 2023). SARS-CoV-2 RNPs appear highly heterogeneous in EM (Carlson et al., 2022; Landeras-Bueno et al., 2025; Yao et al., 2020), and this is reflected in the polymorphic oligomeric states of RNP species we observe in SV-AUC and MP, that we believe is driven by promiscuous self-association or clustering of transient LRS helices (Zhao et al., 2022). Extending previously described characteristics of fuzziness in protein complexes (Duro et al., 2015; Fuxreiter, 2018; Tompa and Fuxreiter, 2008), plasticity seems to involve even basic architectural principles, considering not only the emergence of new distant stabilizing interfaces such as described here in the N-arm, but also the possibility of RNP assembly of truncated N_210-419_* lacking one of the major nucleic acid binding interfaces (Adly et al., 2023; Bouhaddou et al., 2023; Mears et al., 2025; Mulloy et al., 2025; Syed et al., 2024) (see below).

Unfortunately, this intrinsic heterogeneity poses significant methodological hurdles. Nonetheless, salient structural features and assembly principles may be derived from constraints of known binding interfaces and oligomeric states of the RNP and its subunits, as observed in SV-AUC and MP. While the arrangement sketched in **Figure 1C** satisfies these requirements, alternate less symmetrical configurations can be conceived that seem at least equally likely and may coexist in polydisperse mixtures of RNPs. For example, there is no evidence to exclude the possibility of anti-parallel LRS helices pointing the folded nucleic acid -binding domains in different relative orientations, or of mixed co-assemblies with N_210-419_* subunits lacking the NTD (**Supplementary Figure S1**). Uniformity of N-protein/RNA clusters may not be relevant for adequate gRNA condensation.

Beyond the structural model, to study the effect of a larger number of N-protein mutations derived from variants of concern side-by-side in the context of virus assembly, we have carried out experiments using a VLP assay (Syed et al., 2021) (**Figure 7**). In these experiments, all four structural proteins are transfected into 293T cells to package a reporter RNA into VLPs and their infection of receiver cells can be compared. While this assay has been widely used for rapid assessment of spike protein and N variants (Syed et al., 2021), it has limitations due to the addition of non-genomic RNA and the lack of double membrane vesicles from which gRNA emerges through the NSP3/NSP4 pore complex potentially poised for packaging (Bessa et al., 2022; Ke et al., 2024; Ni et al., 2023). It should also be recognized that the results do not directly reflect the relative efficiency of RNP assembly only, since protein expression levels, their localization, and their posttranslational modifications are not controlled for. Susceptibility for such factors might be exacerbated with mutations that modulate weak protein interactions. For example, as shown previously (Syed et al., 2024; Zhao et al., 2024), a GSK3 inhibitor inhibiting N-protein phosphorylation significantly enhances VLP formation and eliminates the advantage provided for by the N:G215C mutation relative to the ancestral N – presumably due to an increase in assembly-competent, non-phosphorylated N-protein erasing an affinity advantage. A similar process may be underlying the absent or marginal improvement in VLP readout from the cysteine LRS mutants and P13L at the achieved transfection level in the present work, and the enhanced signal from R203K/G204R and R203M (the latter being consistent with previous reports (Li et al., 2025; Syed et al., 2021)) modulating protein phosphorylation. Nonetheless, mirroring the results of the biophysical *in vitro* experiments, the addition of RNP-stabilizing P13L and G214C mutations on top of R203K/G204R led to a significantly larger VLP signal.

The VLP assay may be limited in sensitivity to mutation effects due to its restriction to a single round of infection. To avoid this and other potential limitations of the VLP assay for the study of viral packaging, for the key mutation N:P13L we carried out reverse genetics experiments. These showed the sole N:P13L mutation significantly increases viral fitness (**Figure 8**).

Regarding the cysteine mutations that have been repeatedly introduced in the LRS prior to the rise of the Omicron variants of concern, it is an open question whether they lead to covalent bonds *in vivo or in the V*LP assay. While examples of disulfide-linked viral nucleocapsid proteins have been reported (Kubinski et al., 2024; Prokudina et al., 2004; Wootton and Yoo, 2003), a methodological difficulty in their detection is artifactual disulfide bond formation post-lysis of infected cells (Kubinski et al., 2024; Wootton and Yoo, 2003). However, our results clearly show that a major effect of the cysteines already arises in reduced conditions without any covalent bonds, through extension of the LRS helices, and concomitant redirection of the disordered N-terminal sequence. While oxidized tetrameric N-proteins of N:G214C and N:G215C can be incorporated into RNPs, the covalent bonds provided only marginally improved RNP stability. Interestingly, the introduction of cysteines imposes preferences of RNP oligomeric states dependent on oxidation state, consistent with our MD simulations highlighting the impact of cysteine orientation of 214C *versus 215C relati*ve to the hydrophobic surface of the LRS helices. Overall, considering potentially detrimental structural constraints from covalent bonds on LRS clusters seeding RNPs, energetic penalties on RNP disassembly, as well as the required monomeric state of the LRS helix for interaction with the NSP3 Ubl domain (Bessa et al., 2022), at present it is unclear to what extent the formation of disulfide linkages between LRS helices would be beneficial or detrimental in the viral life cycle.

Recent work by the Soranno laboratory has identified an additional function of the disordered N-arm in transiently interacting with the NTD (Cubuk et al., 2021) and dynamically enhancing the affinity of the NTD for RNA (Cubuk et al., 2024). Using single-molecule Förster Resonance Energy Transfer (smFRET) a fourfold modulatory effect of the P13L/Δ31-33 mutation on the NTD RNA binding affinity was observed in N-arm-NTD constructs. Through MD simulations the reduced NTD affinity for RNA was attributed to the N-arm Δ31-33 deletion (Cubuk et al., 2024). Superficially, this may seem in slight conflict with our results of similar T_10_ affinity of full-length ancestral N with and without the P13L/Δ31-33 mutation, but results were obtained in different buffer conditions (50 mM TRIS, pH 7.4 in (Cubuk et al., 2024) *versus 20 mM HEPES*, 150 mM NaCl, pH 7.5 in the present work). In any event, RNA binding of NTD and stabilization of the RNP are different processes; any modulation of N-arm contributions to NTD-RNA interactions through Omicron N-arm mutations Δ31-33 may coexist and be over-compensated for by N-arm self-association interfaces through P13L modulating RNP subunit interactions in the high local N-arm density of the RNP.

The double mutant R203K/G204R arose early in the pandemic and was adopted in several variants of concern (including Alpha, Gamma, Lambda, Zeta, and Omicron BA.1) with the triple nucleotide changes G28881A, G28882A, and G28883C (Mears et al., 2025; Syed et al., 2024) (**Supplementary Figure S12**). As mentioned above, on the protein level N:R203K/G204R has been shown to alter phosphorylation (though in different ways in *in vitro VLP or in vivo* reverse genetics experiments) (Johnson et al., 2022; Syed et al., 2024; Yun et al., 2022); and phosphorylation in turn reduces nucleic acid binding and promotes viral replication as opposed to assembly functions (Botova et al., 2024; Bouhaddou et al., 2023; Carlson et al., 2020; Syed et al., 2024). Adding to such a switch, in the present work we observed the loss of RNP stability of N:R203K/G204R relative to the ancestral N, extending the previous observation of reduced LLPS propensity of N:R203K/G204R (Nguyen et al., 2024). Simultaneously, on the RNA level the N:R203K/G204R mutations also lead to the new formation of a TRS sequence ACGAAC underlying the expression of N_210-419_* in virus-infected cells (though not expected to occur with N:R203K/G204R in the VLP assay lacking the viral RNA-dependent RNA polymerase). It has been hypothesized that N_210-419_* confers increased viral fitness through the suppression of the host anti-viral response (Mears et al., 2025; Mulloy et al., 2025), and that it can assist RNP formation (Bouhaddou et al., 2023; Syed et al., 2024). However, the contribution of N_210-419_* to assembly is still unclear: although it is remarkably capable of forming RNPs *in vitro* and VLP assays (Adly et al., 2023; Bouhaddou et al., 2023; Syed et al., 2024), in infected cells and virions N_210-419_* has been detected only as a minority species (Mears et al., 2025; Mulloy et al., 2025). Also, the recent major Omicron XEC variant (Scarpa et al., 2025) (which had close to 60% global frequency at the beginning of 2025; **Supplementary Figure S12**) exhibits a fourth consecutive nucleotide change G28884C that maintains a similar RG mutation forming R203K/G204P but ablates the canonical TRS sequence, such that continued expression of N_210-419_* in XEC is in question. We propose that an alternative or additional mechanism to retain viral assembly functions may be presented by the accompanying P13L mutation, which our data suggest can more than restore loss of RNP stability in the combination of RG mutations with P13L. This combination occurs in N_o_ and all Omicron variants so far, and was even further stabilized with a cysteine in the LRS in N_λ_.

In conclusion, it has been proposed that mutations in SARS-CoV-2 N protein that affect viral assembly can impact infectivity and fitness (Bouhaddou et al., 2023; Syed et al., 2024; Wu et al., 2021; Zhao et al., 2022). We believe the observed modulations of the RNP assembly and stability studied here highlight a key mechanism for this. Although effects on fitness of viruses carrying N mutations are most likely multi-factorial, they have been observed in reverse genetics tissue culture experiments previously for N:R203K/G204R (Johnson et al., 2022; Mears et al., 2025; Wu et al., 2021), N:G215C (Kubinski et al., 2024), and in the present work for N:P13L. On the other hand, the rise of new variants of concern was usually dominated by their spike protein mutations (with the exception of 21I replacement by 21J which has identical spike mutations but acquired N:G215C (Marchitelli et al., 2021; Stern et al., 2021; Zhao et al., 2022) in the rise of Delta variant), and in many cases N mutations of previously dominant variants were completely replaced by another set of N mutations (dramatically exemplified in the displacement of Delta by Omicron variants). This reinforces the view that these N mutations are secondary to alterations in the immune landscape and transmissibility as the primary driver of evolution (Markov et al., 2023). Nonetheless, the remarkable plasticity of RNPs offers multiple avenues to modulate stability and to compensate for potentially RNP-destabilizing effects of mutations that are beneficial in other ways. In convergent evolution, this has been a constant theme of N protein mutations throughout the SARS-CoV-2 pandemics up until to date. We hypothesize that the ‘fuzziness’ and pleomorphic ability of RNP assembly, with its variable distribution of overall binding energy into several different weak or ultra-weak protein interfaces, and the poor structural definition ranging from flexible chain configurations to polydisperse oligomeric states, provides an evolutionary advantage of orchestrated disorder to promote epistatic interactions and facilitate host adaptation.

## Methods

### Protein expression and purification

Full-length N-protein of the wild type and mutant SARS-CoV-2 were expressed and purified as described previously (Nguyen et al., 2024; Zhao et al., 2024) . Briefly, One Shot BL21(DE3)pLysS E.coli (Thermo Fisher Scientific, Carlsbad, CA) was transformed using a pET29a(+) plasmid vector, which contains a kanamycin-resistant gene and the gene encoding the N-protein of interest preceded by 6xHis tag followed by a Tobacco Etch Virus (TEV) cleavage site at the N-terminus. After cell lysis, the protein was purified by Ni^2+^ affinity chromatography, where on-column unfolding by urea and refolding steps were carried out to remove protein-bound bacterial nucleic acid (Carlson et al., 2020). After tag cleavage by TEV protease, 6xHis tag removal was verified through another round of affinity chromatography, and/or via mass spectrometry. Cleaved protein was subjected to a final size exclusion chromatography followed by dialysis into working buffer (20 mM HEPES, 75 mM NaCl, pH 7.50, supplemented with 1 mM TCEP for cysteine containing proteins, unless otherwise mentioned). Protein purity was validated by SDS-PAGE and the absence of nucleic acid was confirmed by an absorbance ratio 260/280 of ≈0.50-0.55. Final protein concentration was determined by UV-Vis spectrophotometry or by refractive index detected SV-AUC.

N peptides were purchased from ABI Scientific (Sterling, VA), purified by HPLC, examined by matrix-assisted laser desorption/ionization for purity and identity, and lyophilized. The oligonucleotide T_10_ and stem–loop RNA SL7 were purchased from Integrated DNA Technologies (Skokie, IL) and purified by HPLC and lyophilized by the vendor. After reconstitution, SL7 was subject to thermal denaturation at 95 °C for 2 min followed by gradual cooling to room temperature over 1–2 h. For sequences of oligonucleotides and peptides see **Supplementary Table S1**.

The 5,5’-Dithiobis-(2-Nitrobenzoic Acid) (DTNB) (catalog #22582) and Cysteine-HCl (catalog #44889) were purchased from Thermo Fisher Scientific Inc. (Waltham, MA). Free thiols in the protein samples were quantified using the Ellman’s assay by following the standard protocol from the vendor. Briefly, free thiols react with DTNB, resulting in a measurable yellow-colored product, TNB. The quantity of sulfhydryl groups was calculated by the absorbance of the sample using the molar extinction coefficient of TNB at 412 nm (14,150 M^-1^cm^-1^). The same samples were analyzed by SDS-PAGE without the addition of a reducing agent. The relative intensities of the monomer and dimer bands reflect the amounts of the reduced and oxidized forms, respectively.

### Structure prediction and MD simulations

In the studies of LRS peptides, the initial structures of the monomer and oligomers of the ancestral N_210-246_ (**Supplementary Table S1**) were predicted using AlphaFold3 (AF3). The predicted conformations agreed with those reported in (Zhao et al., 2023, 2022), with the oligomers showing a parallel, left-handed coiled-coil signature. Point mutations were introduced by replacing the corresponding glycine residue with cysteine. Graphics were created using ChimeraX.

MD simulations were performed for monomers and trimers of the ancestral peptides N_210-246_ and two mutant sequences (G214C and G215C), following the protocol described in (Zhao et al., 2023). In each case, the initial structure was the top-ranked AF3 model. Each simulation was extended for 250 ns after thermal equilibration under experimental conditions (T = 20°C, P = 1 atm, pH 7.5, and 75 mM NaCl) using the isothermal-isobaric ensemble as implemented in NAMD. The all-atom representation of the CHARMM (param36) force field was used. Structural stabilization of the helical regions was observed within a few nanoseconds, and data were analyzed over the last 200 ns of the simulations.

Structure of the N-arm oligomers was predicted using ColabFold (Mirdita et al., 2022), assembling 20 copies of ancestral N_10-20_, N_10-20_:P13L, N_10-20_:P13S, or N_10-20_:P13T. Structures were analyzed and displayed using ChimeraX (Pettersen et al., 2021).

### Electron microscopy

Carbon coated 200 mesh copper TEM grids were glow discharged for ≈15 seconds. Next, 4 µL of the sample solution was deposited onto the grids and incubated for 2 minutes. After incubation, excess sample solution was removed by gently touching the edge of the grids with filter paper. A large drop of distilled water was then placed on the grids for 1 minute, followed by removal of the water by contacting the grid edge with filter paper. This rinsing step was repeated three times. The grids were then stained with 5 µL of 1% uranyl acetate (UA) solution for 20 seconds. Any excess UA was removed by touching the edge of the grids with filter paper and then left to air dry. Finally, the grids were examined with a FEI Tecnai12 Transmission Electron Microscope (FEI, Hillsboro, Oregon), operating at 120 keV beam energy. TEM images are captured using a high-speed, high-resolution Gatan Rio 3k x 3k CMOS camera (Gatan, Warrendale, PA).

### Sedimentation velocity analytical ultracentrifugation

SV-AUC experiments were conducted in a ProteomeLab XL-I analytical ultracentrifuge (Beckman Coulter, Indianapolis, IN) using standard protocols as previously described (Schuck et al., 2015). AUC cell assemblies filled with samples using 12- or 3-mm charcoal-filled Epon double-sector centerpieces with sapphire windows were loaded into An-50 or An-60 rotors and temperature equilibrated in the rotor chamber at 20 °C for 2-3 hrs. Subsequently, radial scans were acquired with Rayleigh interference optics and absorbance optics at 260 nm and/or 280 nm. Calibration factors for the instrument were determined according to previously published methods (Ghirlando et al., 2013). Sedimentation boundary data were analyzed using sedimentation coefficient distribution c(s) model in the software SEDFIT (Schuck, 2016) (https://sedfitsedphat.nibib.nih.gov/software).

### Mass photometry

Mass photometry measurements were performed in a TwoMP instrument (Refeyn, UK) following the standard protocol (Wu and Piszczek, 2021) unless otherwise mentioned. Samples were loaded in the mini-wells formed by a silicone gasket which was placed on top of a coverslip mounted on the microscope stage. Two configurations of sample loading were used in the current study. For the time-dependent experiments, the samples were first prepared in the Eppendorf tubes prior to MP experiment. Then the working buffer (9 μL) was loaded onto the coverslip for focusing the objective.

Subsequently, the sample was added to the buffer droplet, gently mixed and the measurement was initiated immediately as the 1^st^ time point. The subsequent acquisitions for the same sample continued for specific time intervals. For the samples which were not subject to this dilution/mixing configuration, sample mixtures were equilibrated 2 – 3 hours, focusing was achieved by using the buffer-free option provided by the data acquisition software, and data was collected immediately after sample application. In either configuration, the impact of surface binding on the sample concentration is < 1% and negligible, as described in the **Supplementary Methods S1**. Calibration of the TwoMP instrument was performed using the two calibrants, Beta-Amylase from Sweet Potato (Sigma A8781) and Thyroglobulin from Bovine Thyroid (Sigma T9145) as recommended by the manufacturer. MP data was acquired with AcquireMP software and the analysis was performed with DiscoverMP software (Refeyn, UK).

Rapid mixing experiments were carried out on a OneMP-MassFluidix HC system (Refeyn Ltd., Oxford, UK). A rapid-dilution microfluidic chip (MP-CON-51001, Refeyn Ltd., Oxford, UK) was connected to the buffer, sample, and waste lines. Sample was injected into the chip at a flow rate of 8 µL/min, while the buffer was flowed at 1100 µL/min, and 1 min videos were recorded for data acquisition.

### Circular dichroism spectroscopy

CD spectra were acquired in a Chirascan Q100 (Applied Photophysics, UK) at 20 °C. Measurements were performed in 0.1 mm (peptides) or 1 mm (proteins) pathlength cuvettes with 1 nm steps, and a 1 sec integration time per data point. Each spectrum represents the average of three independent scans with background subtraction applied.

### Dynamic light scattering

Dynamic light scattering measurements of the samples were performed in a Prometheus Panta (Nanotemper, Germany) instrument at 20°C. The samples were loaded into capillaries (Nanotemper PR-AC002) and autocorrelation functions (ACFs) were acquired using the 405 nm laser at the detection angle of 140°. The ACFs were analyzed with discrete species models, size-distribution models, and cumulant analysis in SEDFIT (Parker and Lollar, 2021).

### Virus-like particle assay

The plasmids pLVX-EF1alpha-SARS-CoV-2-E-2xStrep-IRES-Puro (#141385), pLVX-EF1alpha-SARS-CoV-2-M-2xStrep-IRES-Puro (#141386), and pLVX-EF1alpha-SARS-CoV-2-N-2xStrep-IRES-Puro (#141391) were obtained from Addgene. Plasmid pLuc-T20 was a kind gift from Jennifer A. Doudna. The plasmid pIRES2-EGFP (Cat # V011106) was purchased from NovoPro. Plasmids encoding ACE2, pcDNA3.1(+)-SARS-CoV-2 WA1-S and pGAGGS-TMPRSS2 were previously described (Shi et al., 2022, 2021). To construct the plasmid pcDNA3.1(+)-N, the N gene was cloned into pcDNA3.1(+) between the BamHI and NotI sites. Mutations in N were generated by QuikChange™ site-directed mutagenesis, and verified by whole plasmid sequencing. Similarly, to prepare plasmid pIRES2-ME, the M gene was first cloned into pIRES2-EGFP between NheI and BamHI sites, followed by the insertion of the E gene into the resulting plasmid between NotI and a PvuI site introduced after the start codon of EGFP.

The SARS-CoV-2 virus-like particles (VLPs) were prepared as previously described(Syed et al., 2021; Zhao et al., 2024) with some modifications. For SC2-VLP production, 0.8×10^6^ 293T cells (CLS Cat# 305117, RRID:CVCL_1926) per well were plated in a 6-well plate and allowed to grow 24 hours before transfection. Plasmids Cov2-N (1.34), CoV2-M-IRES-E (0.659), CoV-2-Spike (0.0032) and Luc-T20 (2.0) at indicated mass ratios for a total of 4 µg of DNA were diluted in 250 µL Opti-MEM reduced serum medium (Gibco cat# 31985062) at room temperature. 12 µL of TransIT®-293 Transfection Reagent (Mirus Bio, cat# MIR 2704) equilibrated to room temperature was added to each sample, and immediately vortexed. The transfection mixtures were incubated for 25 min at room temperature and then added dropwise to 293T cells after media change with 2 mL of DMEM containing fetal bovine serum and penicillin/streptomycin. Media was changed after 24 h of transfection and at 48 h post-transfection, VLP containing supernatant was collected and filtered using a 0.22 µm syringe filter. Cells in each well were rinsed with 1 mL of PBS and then lysed directly in the wells with 300 µL of NuPAGE™ LDS Sample Buffer (Invitrogen, cat# NP0008) containing HALT protease inhibitors (Thermo Scientific, cat# 87786) and 10 mM DTT.

For the luciferase assay, the wells in 12-well plates were pre-treated with 1 mL poly-D-Lysine (Thermo Fisher Scientific, cat# A3890401) for 30 min, which was removed, washed with PBS once, and finally the plates were airdried. In each well of these pre-treated plates, 1.5×10^5^ receiver cells (293T cells transfected with the plasmids encoding ACE2 and pGAGGS-TMPRSS2 at a mass ratio of 1:1 using the TransIT®-293 transfection reagent) were plated. Next day, the media was replaced, and the cells infected with 250 µL of supernatant from the producer cells. After 24 h, the media was removed, and cells were rinsed with PBS and lysed in 150 µL passive lysis buffer (Luciferase Assay System, Promega, cat# E1500) for 15 min at room temperature with gentle rocking. 20 µL of each lysate was transferred to an opaque black 96-well plate in triplicate, and 50 µL of reconstituted luciferase assay buffer was added and mixed with each lysate. Luminescence was measured immediately after mixing using a TECAN plate reader.

### Generation of recombinant SARS-CoV-2

The generation and use of recombinant SARS-CoV-2 (rSARS-CoV-2) viruses in tissue culture at biosafety level 3 were approved by NIH Institutional Biosafety (IBC) and the Dual Use Research of Concern Institutional Review Entity (DURC-IRE) Committees (IBC approved case number: RD-22-XI-11).

We generated rSARS-CoV-2 viruses using a bacterial artificial chromosome (BAC)-based SARS-CoV-2 reverse genetics system (C. Ye et al., 2020), with detailed construction and recovery procedures described in our previous study (T. Li et al., 2023). Specifically, we fused the mCherry gene with the N gene via a 2A linker and created an intermediate plasmid, pUC57-NEM, containing a portion of the pBAC-SARS-CoV-2 genome digested with BamHI and SalI restriction enzymes. The P13L mutation was introduced via site-directed mutagenesis on the pUC57-NEM plasmid. PCR amplification was performed to generate fragments containing the NEM genes, which were then assembled with the BamHI/SalI-digested larger fragment using the NEBuilder® HiFi DNA Assembly Master Mix. A furin cleavage site mutation (R685S) served as the backbone for introducing mutations in the N protein.

Plasmids were purified using the QIAGEN Plasmid Maxi Kit. Confluent BHK21-ACE2 cells (2 × 10^6^ cells/well in 6-well plates, in duplicate, RRID: CVCL_1914, ATCC) were transfected with 2.5 μg/well of pBAC-SARS-CoV-2 using the TransIT-LT1 transfection reagent. After 6 hours, the medium was replaced with fresh DMEM containing 2% FBS. At 48-72 hours post-transfection, mCherry-positive cells exhibiting signs of viral infection were detached, collected with the supernatant, labeled as P0, and stored at−80°C. The P0 viral stock was centrifuged to remove cell debris and used to infect fresh Vero E6-TMPRSS2 (RRID: CVCL_0574, ATCC) cells for 48-72 hours. The resulting P1 viral stock was collected, and viral titers were determined following next-generation sequencing (NGS) confirmation of the viral genome sequence.

### SARS-CoV-2 infection

Vero E6-TMPRSS2 and A549-hACE2 (RRID: CVCL_0023, ATCC) cell lines were seeded in 12-well plates at 3 × 10^5^ cells/well. The next day, cells were infected at MOI 0.01 with an inoculation period of 1 hour, followed by a medium change to fresh culture medium. Supernatants were collected, and infected cells were fixed with 4% paraformaldehyde (PFA) for 30 minutes at indicated time before removal from the BSL-3 laboratory for fluorescence imaging using Cytation 5 (BioTek). Virus released into the supernatant was titrated using Vero cells via the limiting dilution method. Cell lines were confirmed to be mycoplasma-free using MycoStrip (InvivoGen, rep-mys-50).

## Acknowledgements

This work was supported by the Intramural Research Programs of NIBIB (ZIA EB000099-02), NHLBI, NCI, and NIAID at the National Institutes of Health (NIH). The contributions of the NIH authors are considered Works of the United States Government. The findings and conclusions presented in this paper are those of the authors and do not necessarily reflect the views of the NIH or the U.S. Department of Health and Human Services. We thank the Biophysics Resource in the Center for Structural Biology, Center for Cancer Research, NCI at Frederick for assistance with LC-MS studies. This work utilized the computational resources of the NIH HPC Biowulf cluster for molecular dynamics simulations.

## References

Adly AN, Bi M, Carlson CR, Syed AM, Ciling A, Doudna JA, Cheng Y, Morgan DO. 2023. Assembly of SARS-CoV-2 ribonucleosomes by truncated N_∗_ variant of the nucleocapsid protein. Journal of Biological Chemistry 299:105362. doi:10.1016/j.jbc.2023.105362

Alberti S, Gladfelter AS, Mittag T. 2019. Considerations and Challenges in Studying Liquid-Liquid Phase Separation and Biomolecular Condensates. Cell 176:419–434. doi:10.1016/j.cell.2018.12.035

Alderson TR, Pritišanac I, Kolarić Đ, Moses AM, Forman-Kay JD. 2023. Systematic identification of conditionally folded intrinsically disordered regions by AlphaFold2. Proceedings of the National Academy of Sciences 120:2022.02.18.481080. doi:10.1073/pnas.2304302120

Alston JJ, Soranno A, Holehouse AS. 2023. Conserved molecular recognition by an intrinsically disordered region in the absence of sequence conservation. bioRxiv. doi:10.1101/2023.08.06.552128

Barozi V, Edkins AL, Tastan Bishop Ö. 2022. Evolutionary progression of collective mutations in Omicron sub-lineages towards efficient RBD-hACE2: Allosteric communications between and within viral and human proteins. Comput Struct Biotechnol J 20:4562–4578. doi:10.1016/j.csbj.2022.08.015

Bessa LM, Guseva S, Camacho-Zarco AR, Salvi N, Maurin D, Perez LM, Botova M, Malki A, Nanao M, Jensen MR, Ruigrok RWH, Blackledge M. 2022. The intrinsically disordered SARS-CoV-2 nucleoprotein in dynamic complex with its viral partner nsp3a. Sci Adv 8. doi:10.1126/sciadv.abm4034

Bloom JD, Neher RA. 2023. Fitness effects of mutations to SARS-CoV-2 proteins. Virus Evol 9:2023.01.30.526314. doi:10.1093/ve/vead055

Botova M, Camacho-Zarco AR, Tognetti J, Bessa LM, Guseva S, Mikkola E, Salvi N, Maurin D, Herrmann T, Blackledge M. 2024. A specific phosphorylation-dependent conformational switch in SARS-CoV-2 nucleocapsid protein inhibits RNA binding. Sci Adv 10:1–15. doi:10.1126/sciadv.aax2323

Bouhaddou M, Reuschl AK, Polacco BJ, Thorne LG, Ummadi MR, Ye C, Rosales R, Pelin A, Batra J, Jang GM, Xu J, Moen JM, Richards AL, Zhou Y, Harjai B, Stevenson E, Rojc A, Ragazzini R, Whelan MVX, Furnon W, De Lorenzo G, Cowton V, Syed AM, Ciling A, Deutsch N, Pirak D, Dowgier G, Mesner D, Turner JL, McGovern BL, Rodriguez ML, Leiva-Rebollo R, Dunham AS, Zhong X, Eckhardt M, Fossati A, Liotta NF, Kehrer T, Cupic A, Rutkowska M, Mena I, Aslam S, Hoffert A, Foussard H, Olwal CO, Huang W, Zwaka T, Pham J, Lyons M, Donohue L, Griffin A, Nugent R, Holden K, Deans R, Aviles P, Lopez-Martin JA, Jimeno JM, Obernier K, Fabius JM, Soucheray M, Hüttenhain R, Jungreis I, Kellis M, Echeverria I, Verba K, Bonfanti P, Beltrao P, Sharan R, Doudna JA, Martinez-Sobrido L, Patel AH, Palmarini M, Miorin L, White K, Swaney DL, Garcia-Sastre A, Jolly C, Zuliani-Alvarez L, Towers GJ, Krogan NJ. 2023. SARS-CoV-2 variants evolve convergent strategies to remodel the host response. Cell 186:4597-4614.e26. doi:10.1016/j.cell.2023.08.026

Brocca S, Grandori R, Longhi S, Uversky V. 2020. Liquid–liquid phase separation by intrinsically disordered protein regions of viruses: Roles in viral life cycle and control of virus–host interactions. Int J Mol Sci 21:1–31. doi:10.3390/ijms21239045

Brown CJ, Johnson AK, Dunker AK, Daughdrill GW. 2011. Evolution and disorder. Curr Opin Struct Biol 21:441–446. doi:10.1016/j.sbi.2011.02.005

Carlson CR, Adly AN, Bi M, Howard CJ, Frost A, Cheng Y, Morgan DO. 2022. Reconstitution of the SARS-CoV-2 ribonucleosome provides insights into genomic RNA packaging and regulation by phosphorylation. Journal of Biological Chemistry 298:102560. doi:10.1016/j.jbc.2022.102560

Carlson CR, Asfaha JB, Ghent CM, Howard CJ, Hartooni N, Safari M, Frankel AD, Morgan DO. 2020. Phosphoregulation of Phase Separation by the SARS-CoV-2 N Protein Suggests a Biophysical Basis for its Dual Functions. Mol Cell 80:1092-1103.e4. doi:10.1016/j.molcel.2020.11.025

Chang CK, Sue SC, Yu TH, Hsieh CM, Tsai CK, Chiang YC, Lee SJ, Hsiao HH, Wu WJ, Chang WL, Lin CH, Huang TH. 2006. Modular organization of SARS coronavirus nucleocapsid protein. J Biomed Sci 13:59–72. doi:10.1007/s11373-005-9035-9

Cubuk J, Alston JJ, Incicco JJ, Holehouse AS, Hall KB, Stuchell-Brereton MD, Soranno A. 2024. The disordered N-terminal tail of SARS-CoV-2 Nucleocapsid protein forms a dynamic complex with RNA. Nucleic Acids Res 52:2609–2624. doi:10.1093/nar/gkad1215

Cubuk J, Alston JJ, Incicco JJ, Singh S, Stuchell-Brereton MD, Ward MD, Zimmerman MI, Vithani N, Griffith D, Wagoner JA, Bowman GR, Hall KB, Soranno A, Holehouse AS. 2021. The SARS-CoV-2 nucleocapsid protein is dynamic, disordered, and phase separates with RNA. Nat Commun 12:1936. doi:10.1038/s41467-021-21953-3

Cubuk J, Incicco JJ, Hall KB, Holehouse AS, Stuchell-Brereton MD, Soranno A. 2025. The dimerization domain of SARS-CoV-2 nucleocapsid protein is partially disordered and forms a dynamic high-affinity dimer. Cell Rep Phys Sci 6:102695. doi:10.1016/j.xcrp.2025.102695

Davey NE, Cyert MS, Moses AM. 2015. Short linear motifs - Ex nihilo evolution of protein regulation Short linear motifs - The unexplored frontier of the eukaryotic proteome. Cell Communication and Signaling 13:9–11. doi:10.1186/s12964-015-0120-z

Davey NE, Travé G, Gibson TJ. 2011. How viruses hijack cell regulation. Trends Biochem Sci 36:159–169. doi:10.1016/j.tibs.2010.10.002

Dhamotharan K, Korn SM, Wacker A, Becker MA, Günther S, Schwalbe H, Schlundt A. 2024. A core network in the SARS-CoV-2 nucleocapsid NTD mediates structural integrity and selective RNA-binding. Nature Communications 15:1–16. doi:10.1038/s41467-024-55024-0

Duro N, Miskei M, Fuxreiter M. 2015. Fuzziness endows viral motif-mimicry. Mol Biosyst 11:2821–2829. doi:10.1039/c5mb00301f

Dyson HJ, Wright PE. 2005. Intrinsically unstructured proteins and their functions. Nat Rev Mol Cell Biol 6:197–208. doi:10.1038/nrm1589

Elbe S, Buckland-Merrett G. 2017. Data, disease and diplomacy: GISAID’s innovative contribution to global health. Global Challenges 1:33–46. doi:10.1002/gch2.1018

Evseev D, Magor KE. 2021. Molecular Evolution of the Influenza A Virus Non-structural Protein 1 in Interspecies Transmission and Adaptation. Front Microbiol 12. doi:10.3389/fmicb.2021.693204

Focosi D, Spezia PG, Maggi F. 2024. Subsequent Waves of Convergent Evolution in SARS-CoV-2 Genes and Proteins. Vaccines (Basel) 12:887. doi:10.3390/vaccines12080887

Fuxreiter M. 2018. Fuzziness in protein interactions—a historical perspective. J Mol Biol 430:2278–2287. doi:10.1016/j.jmb.2018.02.015

Ghirlando R, Balbo A, Piszczek G, Brown PH, Lewis MS, Brautigam CA, Schuck P, Zhao H. 2013. Improving the thermal, radial, and temporal accuracy of the analytical ultracentrifuge through external references. Anal Biochem 440:81–95. doi:10.1016/j.ab.2013.05.011

Gupta MN, Uversky VN. 2024. Protein structure–function continuum model: Emerging nexuses between specificity, evolution, and structure. Protein Science 33:1–36. doi:10.1002/pro.4968

Guseva S, Perez LM, Camacho-Zarco A, Bessa LM, Salvi N, Malki A, Maurin D, Blackledge M. 2021. 1H, 13C and 15N Backbone chemical shift assignments of the n-terminal and central intrinsically disordered domains of SARS-CoV-2 nucleoprotein. Biomol NMR Assign 15:255–260. doi:10.1007/s12104-021-10014-x

Hadfield J, Megill C, Bell SM, Huddleston J, Potter B, Callender C, Sagulenko P, Bedford T, Neher RA. 2018. NextStrain: Real-time tracking of pathogen evolution. Bioinformatics 34:4121–4123. doi:10.1093/bioinformatics/bty407

Hagai T, Azia A, Babu MM, Andino R. 2014. Use of Host-like Peptide Motifs in Viral Proteins Is a Prevalent Strategy in Host-Virus Interactions. Cell Rep 7:1729–1739. doi:10.1016/j.celrep.2014.04.052

Holehouse AS, Kragelund BB. 2024. The molecular basis for cellular function of intrinsically disordered protein regions. Nat Rev Mol Cell Biol 25:187–211. doi:10.1038/s41580-023-00673-0

Iserman C, Roden CA, Boerneke MA, Sealfon RSG, McLaughlin GA, Jungreis I, Fritch EJ, Hou YJ, Ekena J, Weidmann CA, Theesfeld CL, Kellis M, Troyanskaya OG, Baric RS, Sheahan TP, Weeks KM, Gladfelter AS. 2020. Genomic RNA Elements Drive Phase Separation of the SARS-CoV-2 Nucleocapsid. Mol Cell 80:1078–1091. doi:10.1016/j.molcel.2020.11.041

Jack A, Ferro LS, Trnka MJ, Wehri E, Nadgir A, Nguyenla X, Fox D, Costa K, Stanley S, Schaletzky J, Yildiz A. 2021. SARS-CoV-2 nucleocapsid protein forms condensates with viral genomic RNA. PLoS Biol 19:e3001425. doi:10.1371/journal.pbio.3001425

Johnson BA, Zhou Y, Lokugamage KG, Vu MN, Bopp N, Crocquet-Valdes PA, Kalveram B, Schindewolf C, Liu Y, Scharton D, Plante JA, Xie X, Aguilar P, Weaver SC, Shi P-Y, Walker DH, Routh AL, Plante KS, Menachery VD. 2022. Nucleocapsid mutations in SARS-CoV-2 augment replication and pathogenesis. PLoS Pathog 18:e1010627. doi:10.1371/journal.ppat.1010627

Ke Z, Zhang H, Wang Y, Wang Jingning, Peng F, Wang Jia, Liu X, Hu H, Li Y. 2024. N-terminus of SARS-CoV-2 nonstructural protein 3 interrupts RNA-driven phase separation of N protein by displacing RNA. Journal of Biological Chemistry 107828. doi:10.1016/j.jbc.2024.107828

Klein S, Cortese M, Winter SL, Wachsmuth-Melm M, Neufeldt CJ, Cerikan B, Stanifer ML, Boulant S, Bartenschlager R, Chlanda P. 2020. SARS-CoV-2 structure and replication characterized by in situ cryo-electron tomography. Nat Commun 11:5885. doi:10.1038/s41467-020-19619-7

Korn SM, Dhamotharan K, Jeffries CM, Schlundt A. 2023. The preference signature of the SARS-CoV-2 Nucleocapsid NTD for its 5’-genomic RNA elements. Nat Commun 14:3331. doi:10.1038/s41467-023-38882-y

Kubinski HC, Despres HW, Johnson BA, Schmidt MM, Estes LK, Pekosz A, Crothers JW, Roychoudhury P, Greninger AL, Jerome KR, Martorelli Di Genova B, Walker DH, Ballif BA, Ladinsky MS, Bjorkman PJ, Menachery VD, Bruce EA. 2024. Variant mutation in SARS-CoV-2 nucleocapsid enhances viral infection via altered genomic encapsidation. bioRxiv 23:e3003115. doi:10.1101/2024.03.08.584120

Landeras-Bueno S, Hariharan C, Avalos RD, Norris AS, Snyder DT, Hastie KM, Harkins S, Zandonatti M, Rajamanickam RR, Olmedillas E, Miller R, Shresta S, Wysocki VH, Saphire EO. 2025. Structural stabilization of the intrinsically disordered SARS-CoV-2 N by binding to RNA sequences engineered from the viral genome fragment. Nat Commun 16. doi:10.1038/s41467-025-61861-4

Li P, Xue B, Schnicker NJ, Wong L-YR, Meyerholz DK, Perlman S. 2023. Nsp3-N interactions are critical for SARS-CoV-2 fitness and virulence. Proceedings of the National Academy of Sciences 120:2017. doi:10.1073/pnas.2305674120

Li T, Kang I, Hu Z, Gibbs J, Ye C, Kosik I, Shi G, Holly J, Kosikova M, Compton A, Martinez-Sobrido L, Johnson RF, Xie H, Yewdell JW. 2023. Syncytia Formation Promotes Virus Resistance to Interferon and Neutralizing Antibodies. bioRxiv. doi:10.1101/2023.12.12.571262

Li Y, Li M, Xiao H, Liao F, Shen M, Ge W, Ou J, Liu Y, Chen L, Zhao Y, Wan P, Liu J, Chen J, Lan X, Wu S, Ding Q, Li G, Zhang Q, Pan P. 2025. The R203M and D377Y mutations of the nucleocapsid protein promote SARS-CoV-2 infectivity by impairing RIG-I-mediated antiviral signaling. PLoS Pathog 21:1– 28. doi:10.1371/journal.ppat.1012886

Longhi S, Bloyet LM, Gianni S, Gerlier D. 2017. How order and disorder within paramyxoviral nucleoproteins and phosphoproteins orchestrate the molecular interplay of transcription and replication. Cellular and Molecular Life Sciences 74:3091–3118. doi:10.1007/s00018-017-2556-3

Marchitelli V, Troise C, Vesuviano O, Parisi A, Zooprofilattico I, Bianco A, Zooprofilattico I, Sambro L Del, Zooprofilattico I, Somma R, Vesuviano O, Ingegneria S, Basilicata U, Natale G De. 2021. Evidence for the dependence of the SARS-Cov-2 Delta high diffusivity on the associated Nlli: G215C nucleocapsid mutation. Res Sq 1–9. doi:10.21203/rs.3.rs-846225/v1

Markov P V, Ghafari M, Beer M, Lythgoe K, Simmonds P, Stilianakis NI, Katzourakis A. 2023. The evolution of SARS-CoV-2. Nat Rev Microbiol 21:361–379. doi:10.1038/s41579-023-00878-2

Mears H V, Young GR, Sanderson T, Harvey R, Barrett-Rodger J, Penn R, Cowton V, Furnon W, De Lorenzo G, Crawford M, Snell DM, Fowler AS, Chakrabarti AM, Hussain S, Gilbride C, Emmott E, Finsterbusch K, Luptak J, Peacock TP, Nicod J, Patel AH, Palmarini M, Wall E, Williams B, Gandhi S, Swanton C, Bauer DLV. 2025. Emergence of SARS-CoV-2 subgenomic RNAs that enhance viral fitness and immune evasion. PLoS Biol 23:1–29. doi:10.1371/journal.pbio.3002982

Mirdita M, Schütze K, Moriwaki Y, Heo L, Ovchinnikov S, Steinegger M. 2022. ColabFold: making protein folding accessible to all. Nat Methods 19:679–682. doi:10.1038/s41592-022-01488-1

Mulloy RP, Evseev D, Bui-Marinos MP, Sharlin N, Corcoran JA. 2025. A truncated SARS-CoV-2 nucleocapsid protein enhances virus fitness by evading antiviral responses. bioRxiv 1–68. doi:10.1101/2025.02.15.638421

Nesmelova I V., Melnikova DL, Ranjan V, Skirda VD. 2019. Translational diffusion of unfolded and intrinsically disordered proteinsProgress in Molecular Biology and Translational Science. Elsevier B.V. pp. 85–108. doi:10.1016/bs.pmbts.2019.05.004

Nguyen A, Zhao H, Myagmarsuren D, Srinivasan S, Wu D, Chen J, Piszczek G, Schuck P. 2024. Modulation of biophysical properties of nucleocapsid protein in the mutant spectrum of SARS-CoV-2. Elife 13. doi:10.7554/eLife.94836

Ni X, Han Y, Zhou R, Zhou Y, Lei J. 2023. Structural insights into ribonucleoprotein dissociation by nucleocapsid protein interacting with non-structural protein 3 in SARS-CoV-2. Commun Biol 6:193. doi:10.1038/s42003-023-04570-2

Obermeyer F, Jankowiak M, Barkas N, Schaffner SF, Pyle JD, Yurkovetskiy L, Bosso M, Park DJ, Babadi M, MacInnis BL, Luban J, Sabeti PC, Lemieux JE. 2022. Analysis of 6.4 million SARS-CoV-2 genomes identifies mutations associated with fitness. Science (1979) 1208:1–14. doi:10.1126/science.abm1208

Olsen JG, Teilum K, Kragelund BB. 2017. Behaviour of intrinsically disordered proteins in protein–protein complexes with an emphasis on fuzziness. Cellular and Molecular Life Sciences 74:3175–3183. doi:10.1007/s00018-017-2560-7

Oulas A, Zanti M, Tomazou M, Zachariou M, Minadakis G, Bourdakou MM, Pavlidis P, Spyrou GM. 2021. Generalized linear models provide a measure of virulence for specific mutations in SARS-cov-2 strains. PLoS One 16:1–17. doi:10.1371/journal.pone.0238665

Paravastu AK, Leapman RD, Yau W-M, Tycko R. 2008. Molecular structural basis for polymorphism in Alzheimer’s β-amyloid fibrils. Proceedings of the National Academy of Sciences 105:18349–18354. doi:10.1073/pnas.0806270105

Parker ET, Lollar P. 2021. Measurement of the translational diffusion coefficient and hydrodynamic radius of proteins by dynamic light scattering. Bio Protoc 11:e4195. doi:10.21769/BioProtoc.4195

Perdikari TM, Murthy AC, Ryan VH, Watters S, Naik MT, Fawzi NL. 2020. SARS-CoV-2 nucleocapsid protein phase-separates with RNA and with human hnRNPs. EMBO J 39:1–35. doi:10.15252/embj.2020106478

Pettersen EF, Goddard TD, Huang CC, Meng EC, Couch GS, Croll TI, Morris JH, Ferrin TE. 2021. UCSF ChimeraX: Structure visualization for researchers, educators, and developers. Protein Science 30:70–82. doi:10.1002/pro.3943

Planchais C, Reyes-Ruiz A, Lacombe R, Zarantonello A, Lecerf M, Revel M, Roumenina LT, Atanasov BP, Mouquet H, Dimitrov JD. 2022. Evolutionary trajectory of receptor binding specificity and promiscuity of the spike protein of SARS-CoV-2. Protein Science 31:1–19. doi:10.1002/pro.4447

Prokudina EN, Semenova NP, Chumakov VM, Rudneva IA. 2004. Transient disulfide bonds formation in conformational maturation of influenza virus nucleocapsid protein (NP). Virus Res 99:169–175. doi:10.1016/j.virusres.2003.11.008

Ranganathan S, Dasmeh P, Furniss S, Shakhnovich E. 2023. Phosphorylation sites are evolutionary checkpoints against liquid-solid transition in protein condensates. Proc Natl Acad Sci U S A 120. doi:10.1073/pnas.2215828120

Ranganathan S, Shakhnovich EI. 2020. Dynamic metastable long-living droplets formed by sticker-spacer proteins. Elife 9:1–25. doi:10.7554/eLife.56159

Ren J, Wang S, Zong Z, Pan T, Liu S, Mao W, Huang H, Yan X, Yang B, He X, Zhou F, Zhang L. 2024. TRIM28-mediated nucleocapsid protein SUMOylation enhances SARS-CoV-2 virulence. Nat Commun 15. doi:10.1038/s41467-023-44502-6

Rochman ND, Wolf YI, Koonin E V. 2022. Molecular adaptations during viral epidemics. EMBO Rep 23:1– 22. doi:10.15252/embr.202255393

Roden CA, Dai Y, Giannetti CA, Seim I, Lee M, Sealfon R, McLaughlin GA, Boerneke MA, Iserman C, Wey SA, Ekena JL, Troyanskaya OG, Weeks KM, You L, Chilkoti A, Gladfelter AS. 2022. Double-stranded RNA drives SARS-CoV-2 nucleocapsid protein to undergo phase separation at specific temperatures. Nucleic Acids Res 50:8168–8192. doi:10.1093/nar/gkac596

Różycki B, Boura E. 2022. Conformational ensemble of the full-length SARS-CoV-2 nucleocapsid (N) protein based on molecular simulations and SAXS data. Biophys Chem 288:106843. doi:10.1016/j.bpc.2022.106843

Savastano A, Ibáñez de Opakua A, Rankovic M, Zweckstetter M. 2020. Nucleocapsid protein of SARS-CoV-2 phase separates into RNA-rich polymerase-containing condensates. Nat Commun 11:6041. doi:10.1038/s41467-020-19843-1

Scarpa F, Branda F, Ceccarelli G, Romano C, Locci C, Pascale N, Azzena I, Fiori PL, Casu M, Pascarella S, Quaranta M, Benvenuto D, Cauda R, Ciccozzi M, Sanna D. 2025. SARS-CoV-2 XEC: A Genome-Based Survey. Microorganisms 13. doi:10.3390/microorganisms13020253

Schiavina M, Pontoriero L, Tagliaferro G, Pierattelli R, Felli IC. 2022. The Role of Disordered Regions in Orchestrating the Properties of Multidomain Proteins: The SARS-CoV-2 Nucleocapsid Protein and Its Interaction with Enoxaparin. Biomolecules 12:1302. doi:10.3390/biom12091302

Schuck P. 2016. Sedimentation Velocity Analytical Ultracentrifugation: Discrete Species and Size-Distributions of Macromolecules and Particles. Boca Raton, FL: CRC Press.

Schuck P. 2010. Sedimentation patterns of rapidly reversible protein interactions. Biophys J 98:2005– 2013. doi:10.1016/j.bpj.2009.12.4336

Schuck P, Zhao H. 2023. Diversity of short linear interaction motifs in SARS-CoV-2 nucleocapsid protein. mBio 14:e02388–23. doi:10.1128/mbio.02388-23

Schuck P, Zhao H. 2017. Sedimentation Velocity Analytical Ultracentrifugation: Interacting Systems. Boca Raton, FL: CRC Press.

Schuck P, Zhao H, Brautigam CA, Ghirlando R. 2015. Basic Principles of Analytical Ultracentrifugation. Boca Raton, FL: CRC Press.

Shi G, Chiramel AI, Li T, Lai KK, Kenney AD, Zani A, Eddy AC, Majdoul S, Zhang L, Dempsey T, Beare PA, Kar S, Yewdell JW, Best SM, Yount JS, Compton AA. 2022. Rapalogs downmodulate intrinsic immunity and promote cell entry of SARS-CoV-2. Journal of Clinical Investigation 132:e160766. doi:10.1172/JCI160766

Shi G, Kenney AD, Kudryashova E, Zani A, Zhang L, Lai KK, Hall-Stoodley L, Robinson RT, Kudryashov DS, Compton AA, Yount JS. 2021. Opposing activities of IFITM proteins in SARS-CoV-2 infection. EMBO J 40:e106501. doi:10.15252/embj.2020106501

Stern A, Fleishon S, Kustin T, Mandelboim M, Erster O, Mendelson E, Mor O, Zuckerman NS, Bucris D, Sofer D, Bar-Ilan D, Geva M, Asraf O, Rechavi G, Glick-Saar E, Rainy N, Weiner C, Sorek-Abramovich R, Yegorov Y, Vishnevsky A, Benveniste-Lekovitz P, Ramzia AH, Chaim AB. 2021. The unique evolutionary dynamics of the SARS-CoV-2 Delta variant Israel Consortium of SARS-CoV-2 sequencing. medRxiv 2021.08.05.21261642.

Syed AM, Ciling A, Chen IP, Carlson CR, Adly AN, Martin HS, Taha TY, Khalid MM, Price N, Bouhaddou M, Ummadi MR, Moen JM, Krogan NJ, Morgan DO, Ott M, Doudna JA. 2024. SARS-CoV-2 evolution balances conflicting roles of N protein phosphorylation. PLoS Pathog 20:e1012741. doi:10.1371/journal.ppat.1012741

Syed AM, Taha TY, Tabata T, Chen IP, Ciling A, Khalid MM, Sreekumar B, Chen P-Y, Hayashi JM, Soczek KM, Ott M, Doudna JA. 2021. Rapid assessment of SARS-CoV-2–evolved variants using virus-like particles. Science (1979) 374:1626–1632. doi:10.1126/science.abl6184

Tarczewska A, Kolonko-Adamska M, Zarębski M, Dobrucki J, Ożyhar A, Greb-Markiewicz B. 2021. The method utilized to purify the SARS-CoV-2 N protein can affect its molecular properties. Int J Biol Macromol 188:391–403. doi:10.1016/j.ijbiomac.2021.08.026

Tokuriki N, Oldfield CJ, Uversky VN, Berezovsky IN, Tawfik DS. 2009. Do viral proteins possess unique biophysical features? Trends Biochem Sci 34:53–59. doi:10.1016/j.tibs.2008.10.009

Tokuriki N, Tawfik DS. 2009. Protein Dynamism and Evolvability. Science (1979) 324:203–207. doi:10.1126/science.1169375

Tompa P, Fuxreiter M. 2008. Fuzzy complexes: polymorphism and structural disorder in protein-protein interactions. Trends Biochem Sci 33:2–8. doi:10.1016/j.tibs.2007.10.003

Tugaeva K V., Sysoev AA, Kapitonova AA, Smith JLR, Zhu P, Cooley RB, Antson AA, Sluchanko NN. 2023. Human 14-3-3 Proteins Site-selectively Bind the Mutational Hotspot Region of SARS-CoV-2 Nucleoprotein Modulating its Phosphoregulation. J Mol Biol 435:167891. doi:10.1016/j.jmb.2022.167891

Welkers MRA, Pawestri HA, Fonville JM, Sampurno OD, Pater M, Holwerda M, Han AX, Russell CA, Jeeninga RE, Setiawaty V, de Jong MD, Eggink D. 2019. Genetic diversity and host adaptation of avian H5N1 influenza viruses during human infection. Emerg Microbes Infect 8:262–271. doi:10.1080/22221751.2019.1575700

Wink PL, Volpato FCZ, Monteiro FL, Willig JB, Zavascki AP, Barth AL, Martins AF. 2022. First identification of SARS-CoV-2 lambda (C.37) variant in Southern Brazil. Infect Control Hosp Epidemiol 43:1996– 1997. doi:10.1017/ice.2021.390

Wootton SK, Yoo D. 2003. Homo-Oligomerization of the Porcine Reproductive and Respiratory Syndrome Virus Nucleocapsid Protein and the Role of Disulfide Linkages. J Virol 77:4546–4557. doi:10.1128/jvi.77.8.4546-4557.2003

Wu D, Piszczek G. 2021. Standard protocol for mass photometry experiments. European Biophysics Journal 50:403–409. doi:10.1007/s00249-021-01513-9

Wu H, Fuxreiter M. 2016. The structure and dynamics of higher-order assemblies: amyloids, signalosomes, and granules. Cell 165:1055–1066. doi:10.1016/j.cell.2016.05.004

Wu H, Xing N, Meng K, Fu B, Xue W, Dong P, Tang W, Xiao Y, Liu G, Luo H, Zhu W, Lin X, Meng G, Zhu Z. 2021. Nucleocapsid mutations R203K/G204R increase the infectivity, fitness, and virulence of SARS-CoV-2. Cell Host Microbe 29:1788-1801.e6. doi:10.1016/j.chom.2021.11.005

Wu W, Cheng Y, Zhou H, Sun C, Zhang S. 2023. The SARS-CoV-2 nucleocapsid protein: its role in the viral life cycle, structure and functions, and use as a potential target in the development of vaccines and diagnostics. Virol J 20:6. doi:10.1186/s12985-023-01968-6

Xue B, Blocquel D, Habchi J, Uversky A V., Kurgan L, Uversky VN, Longhi S. 2014. Structural disorder in viral proteins. Chem Rev 114:6880–6911. doi:10.1021/cr4005692

Yao H, Song Y, Chen Y, Wu N, Xu J, Sun C, Zhang J, Weng T, Zhang Z, Wu Z, Cheng L, Shi D, Lu X, Lei J, Crispin M, Shi Y, Li L, Li S. 2020. Molecular Architecture of the SARS-CoV-2 Virus. Cell 183:730-738.e13. doi:10.1016/j.cell.2020.09.018

Ye C, Chiem K, Park JG, Oladunni F, Platt RN, Anderson T, Almazan F, de la Torre JC, Martinez-Sobrido L. 2020. Rescue of SARS-CoV-2 from a single bacterial artificial chromosome. mBio 11:1–10. doi:10.1128/mBio.02168-20

Ye Q, West AMV, Silletti S, Corbett KD. 2020. Architecture and self-assembly of the SARS-CoV-2 nucleocapsid protein. Protein Science 29:1890–1901. doi:10.1002/pro.3909

Yu IM, Gustafson CLT, Diao J, Burgner JW, Li Z, Zhang J, Chen J. 2005. Recombinant severe acute respiratory syndrome (SARS) coronavirus nucleocapsid protein forms a dimer through its C-terminal domain. Journal of Biological Chemistry 280:23280–23286. doi:10.1074/jbc.M501015200

Yun JS, Song H, Kim NH, Cha SY, Hwang KH, Lee JE, Jeong CH, Song SH, Kim S, Cho ES, Kim HS, Yook JI. 2022. Glycogen Synthase Kinase-3 Interaction Domain Enhances Phosphorylation of SARS-CoV-2 Nucleocapsid Protein. Mol Cells 45:911–922. doi:10.14348/molcells.2022.0130

Zachrdla M, Savastano A, Ibáñez de Opakua A, Cima-Omori M, Zweckstetter M. 2022. Contributions of the N-terminal intrinsically disordered region of the severe acute respiratory syndrome coronavirus 2 nucleocapsid protein to RNA-induced phase separation. Protein Science 31:2003–2005. doi:10.1002/pro.4409

Zarin T, Strome B, Peng G, Pritišanac I, Forman-Kay JD, Moses AM. 2021. Identifying molecular features that are associated with biological function of intrinsically disordered protein regions. Elife 10:1– 36. doi:10.7554/eLife.60220

Zhao H, Nguyen A, Wu D, Li Y, Hassan SA, Chen J, Shroff H, Piszczek G, Schuck P. 2022. Plasticity in structure and assembly of SARS-CoV-2 nucleocapsid protein. PNAS Nexus 1:pgac049. doi:10.1093/pnasnexus/pgac049

Zhao H, Syed AM, Khalid MM, Nguyen A, Ciling A, Wu D, Yau W, Srinivasan S, Esposito D, Doudna JA, Piszczek G, Ott M, Schuck P. 2024. Assembly of SARS-CoV-2 nucleocapsid protein with nucleic acid. Nucleic Acids Res 52:6647–6661. doi:10.1093/nar/gkae256

Zhao H, Wu D, Hassan SA, Nguyen A, Chen J, Piszczek G, Schuck P. 2023. A conserved oligomerization domain in the disordered linker of coronavirus nucleocapsid proteins. Sci Adv 9:eadg6473. doi:10.1126/sciadv.adg6473

Zhao H, Wu D, Nguyen A, Li Y, Adão RC, Valkov E, Patterson GH, Piszczek G, Schuck P. 2021. Energetic and structural features of SARS-CoV-2 N-protein co-assemblies with nucleic acids. iScience 24:102523. doi:10.1016/j.isci.2021.102523

Zinzula L, Basquin J, Bohn S, Beck F, Klumpe S, Pfeifer G, Nagy I, Bracher A, Hartl FU, Baumeister W. 2021. High-resolution structure and biophysical characterization of the nucleocapsid phosphoprotein dimerization domain from the Covid-19 severe acute respiratory syndrome coronavirus 2. Biochem Biophys Res Commun 538:54–62. doi:10.1016/j.bbrc.2020.09.131

