## Supplemental Figure S6 for "Evolution of a fuzzy ribonucleoprotein complex in viral assembly"

**Supplementary Figure S6: Structural and physicochemical effects of LRS mutants G214C and G215C under reducing conditions.** Snapshots taken at equal time intervals (4 ns) from the 200-ns MD simulations of the trimeric complex of the ancestral (WT) peptide $N_{210-246}$ and single-point G→C mutants (trimers oriented by overlaying the helical regions of the three monomers; the panels are shown in orthosteric view for ease of comparison). The upper row presents a view from the N-terminus side (compare with Figure 3). The second row provides a side view, showing the tightly packed hydrophobic core that drives and stabilizes monomer self-association. The conformational changes and reorientation of the N-terminal segment observed in the mutant monomers (see Figure 3) influence the interactions between the helices and affect the complex structure and stabilization (see Text). A consequence of this rearrangement is shown in the third row, where E216 (the oxygen atoms of the carboxyl group depicted as red van der Waals spheres), expected to be anionic under the experimental conditions (neutral pH; mimicked in the simulations; see Methods), becomes more spatially concentrated at the helix end. The varying local electric fields (less negative in the WT, slightly more negative in the G214C mutant, and much more negative in the G215C), illustrated on the surface electrostatic potential in the fourth row, may alter, among other features, the binding of cations in the solution, potentially influencing further aggregation of the peptide or full-length N-protein, phase separation, or particle assembly in vivo. Oligomerization under oxidative conditions (not investigated in this study) may differ significantly, as the hydrophobic moieties that drive wild-type helix dimerization could reorient outward in disulfide-bridged dimers, thereby affecting the oligomerization mechanism and the complex's properties.

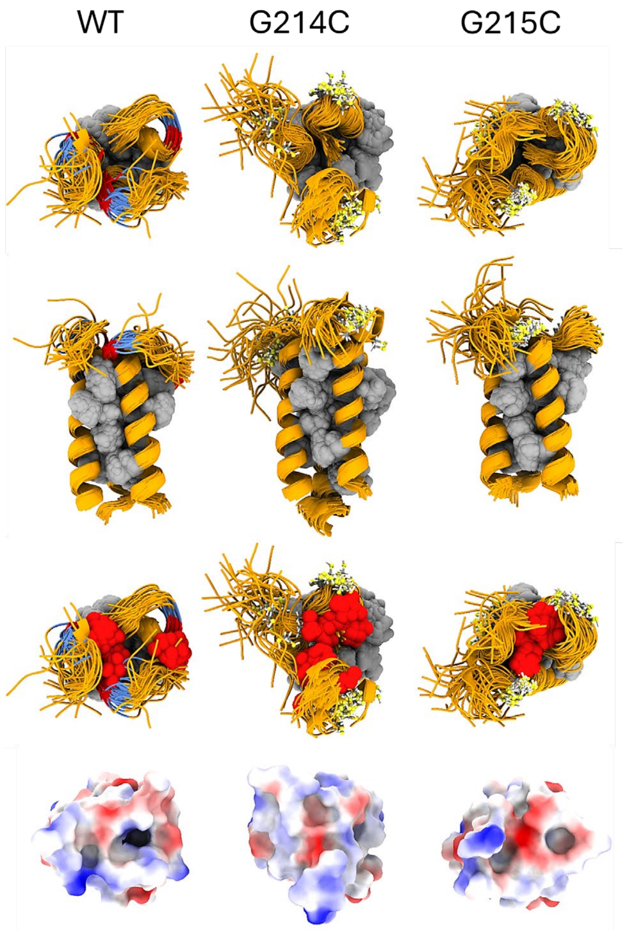
