## Supplemental Figure S7 for "Evolution of a fuzzy ribonucleoprotein complex in viral assembly"

**Supplementary Figure S7: Non-reducing SDS-PAGE of reduced and oxidized N:G215C* and N$_\lambda$*.** Lanes: 1) oxidized N$_\lambda$*; 2) reduced N$_\lambda$; 3) oxidized N:G215C*; 4) reduced N:G215C.

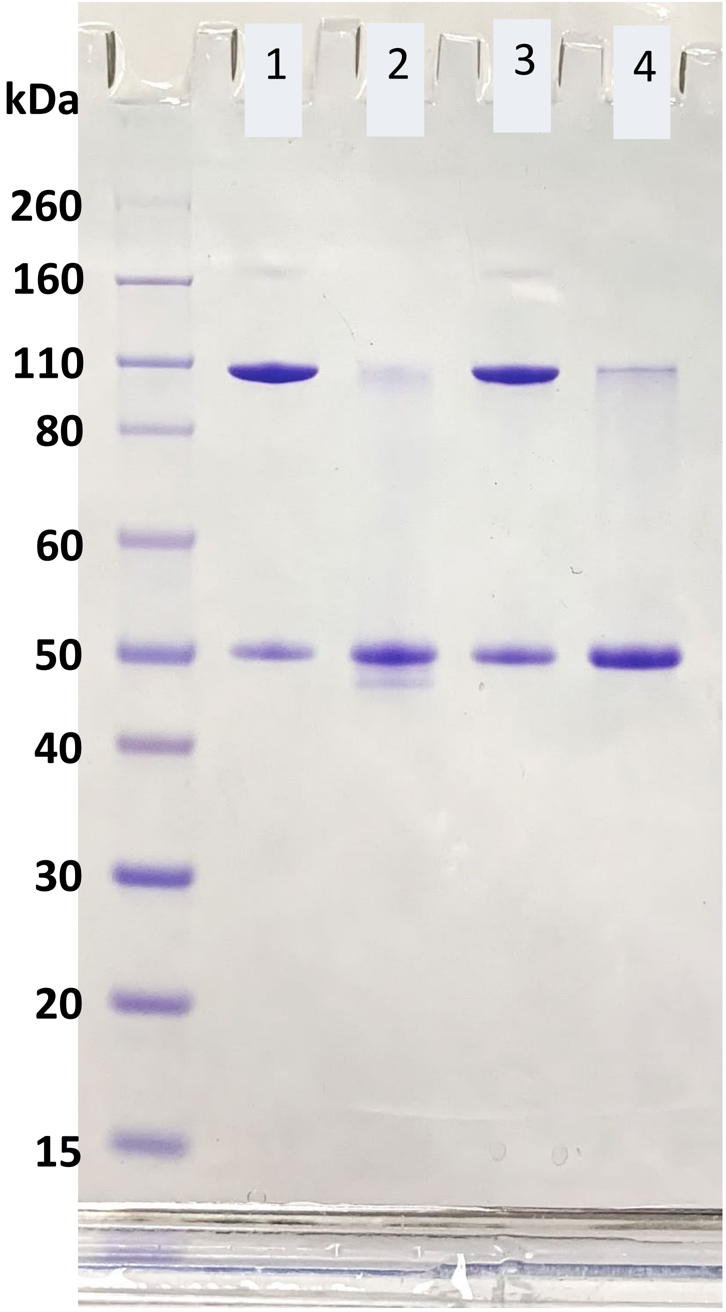
