## Supplemental Figure S8 for "Evolution of a fuzzy ribonucleoprotein complex in viral assembly"

**Supplementary Figure S8: Measurement of the affinity of N:P13L,Δ31-33 for oligonucleotide $T_{10}$ by SV-AUC titration.** Sedimentation coefficient distributions are shown for 1.5 or 3.0 μM protein with $T_{10}$ in different molar ratios. Integration of the overall weight-average *s*-value leads to the isotherm shown in the inset (circles), which is globally fitted with a binding model (lines) resulting in a $K_D$ of 0.88 (0.70 – 1.1) μM. Experiments are in 20mM HEPES, 150mM NaCl, pH 7.5. In comparison, for ancestral N, the best-fit $K_D$-values and 95% confidence intervals are 1.1 [0.8–1.6] μM (Nguyen et al., 2024).

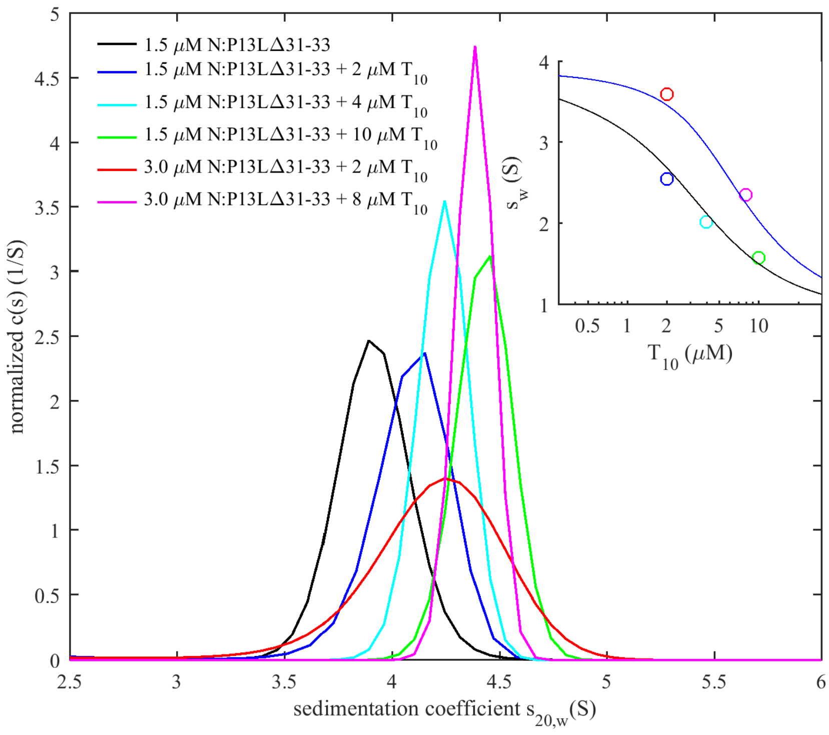
