## Supplemental Figure S9 for "Evolution of a fuzzy ribonucleoprotein complex in viral assembly"

**Supplementary Figure S9: Mutations of N:P13 across the phylogenetic tree of SARS-CoV-2**. Shown are all-time global sequence samples with clade labels and color-coded amino acid at position 13, with the ancestral P13 in green and P13L in yellow. The blue arrow points to the Lambda sequences. Additionally, a cluster of P13L mutations occurred in India in clade 19A. The phylogenetic tree was generated by Nextstrain (Hadfield et al., 2018).

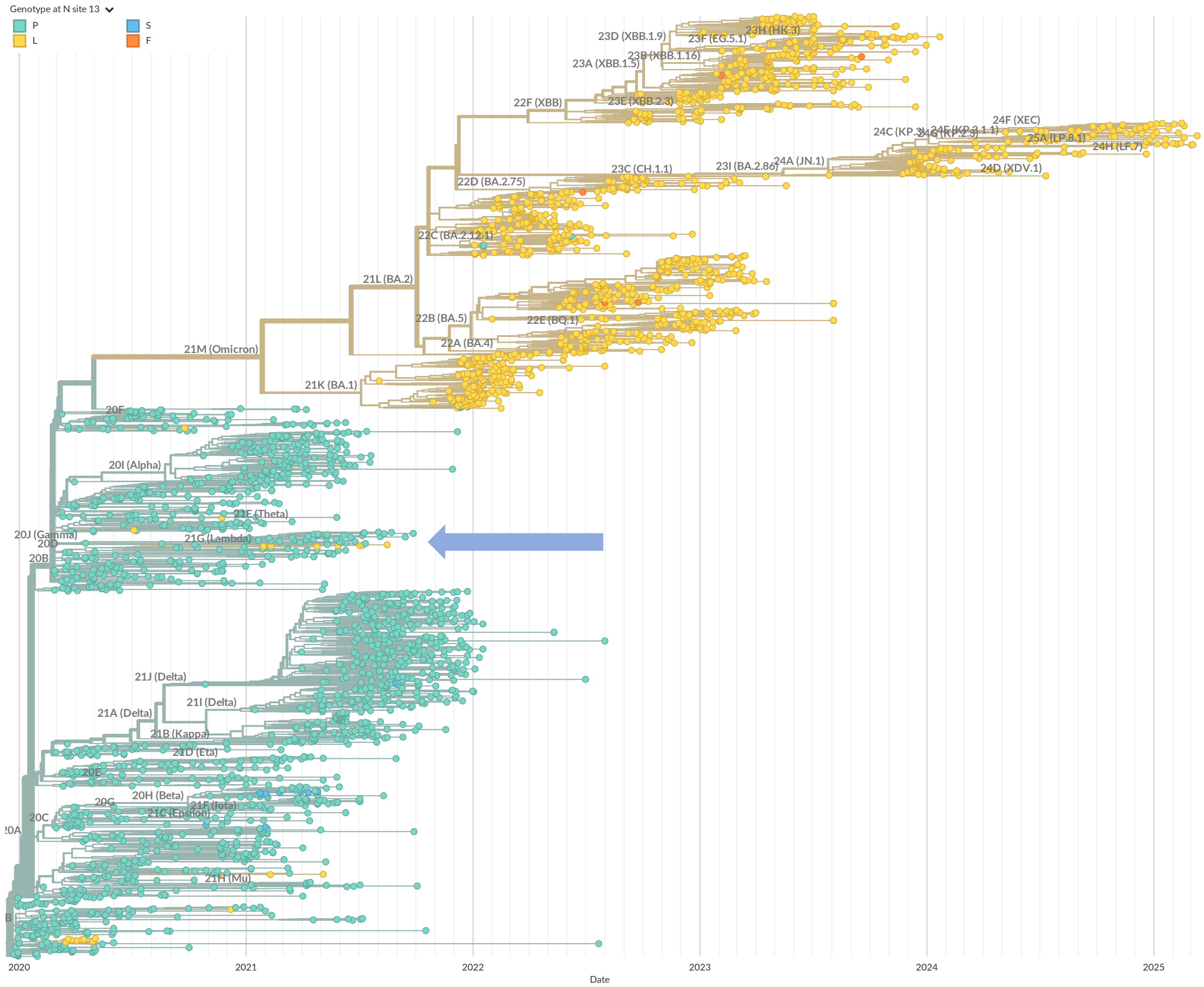
