## Supplemental Figure S10 for "Evolution of a fuzzy ribonucleoprotein complex in viral assembly"

**Supplementary Figure S10: Mutations of N:G215 across the phylogenetic tree of SARS-CoV-2**. Shown are all-time global sequence samples with clade labels and color-coded amino acid at position 215, with the ancestral G215 in green and G215C in yellow. The phylogenetic tree was generated by Nextstrain (Hadfield et al., 2018).

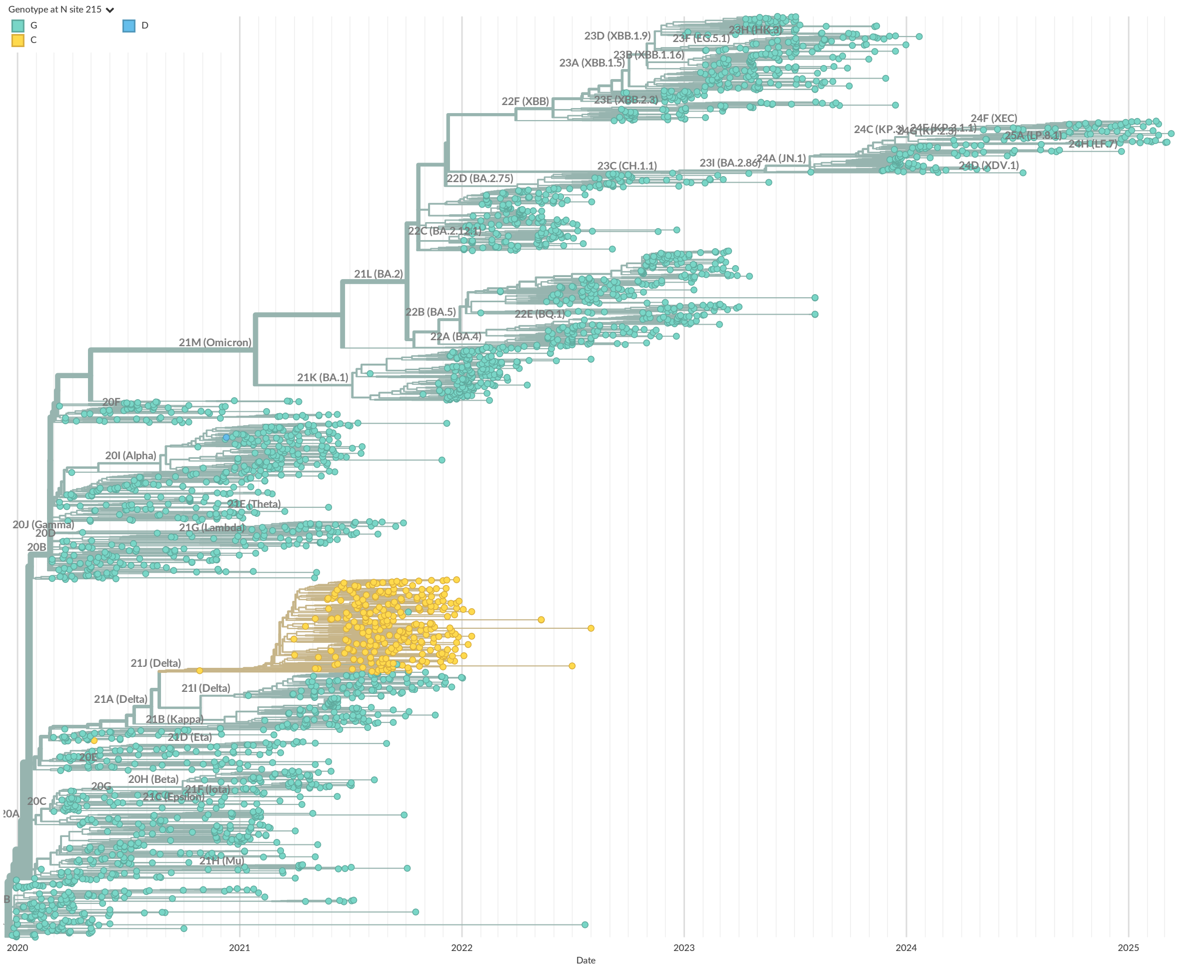
