## Supplemental Figure S11 for "Evolution of a fuzzy ribonucleoprotein complex in viral assembly"

**Supplementary Figure S11: Mutations of N:G214 and N:G215 across the phylogenetic tree of SARS-CoV-2**. Shown are all-time sequence samples in South America with clade labels and color-coded amino acid at position 214 and 215. The combination of 214C/G215 strain 21G (Lambda) is shown in blue, whereas the combination G214/215C of strain 21J (Delta) is shown in yellow. The phylogenetic tree was generated by Nextstrain (Hadfield et al., 2018).

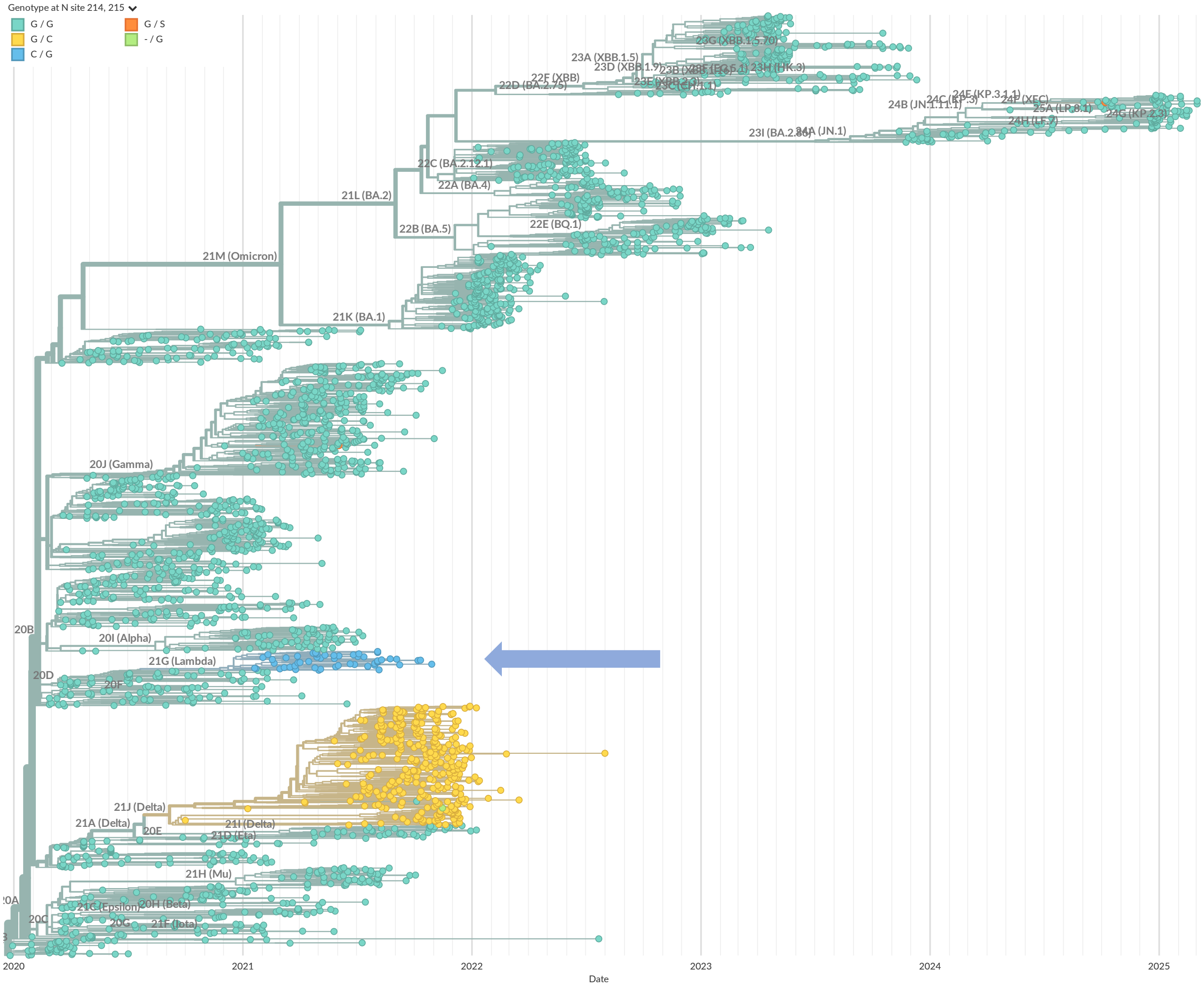
