## Supplemental Figure S12 for "Evolution of a fuzzy ribonucleoprotein complex in viral assembly"

**Supplementary Figure S12: Mutations of N:R203 and N:G204 across the phylogenetic tree of SARS-CoV-2**. Shown are global sequence samples mostly representing sequences of the recent 6 month, with clade labels and color-coded amino acid at position 203 and 204. The ancestral combination of R203/G204 is shown in green, the mutation 203M of the Delta VOC in blue, the combination 203K/204R common to Alpha and Omicron VOCs in yellow, and the combination 203K/204P defining in the Omicron XEC variant in orange. The phylogenetic tree was generated by Nextstrain (Hadfield et al., 2018).

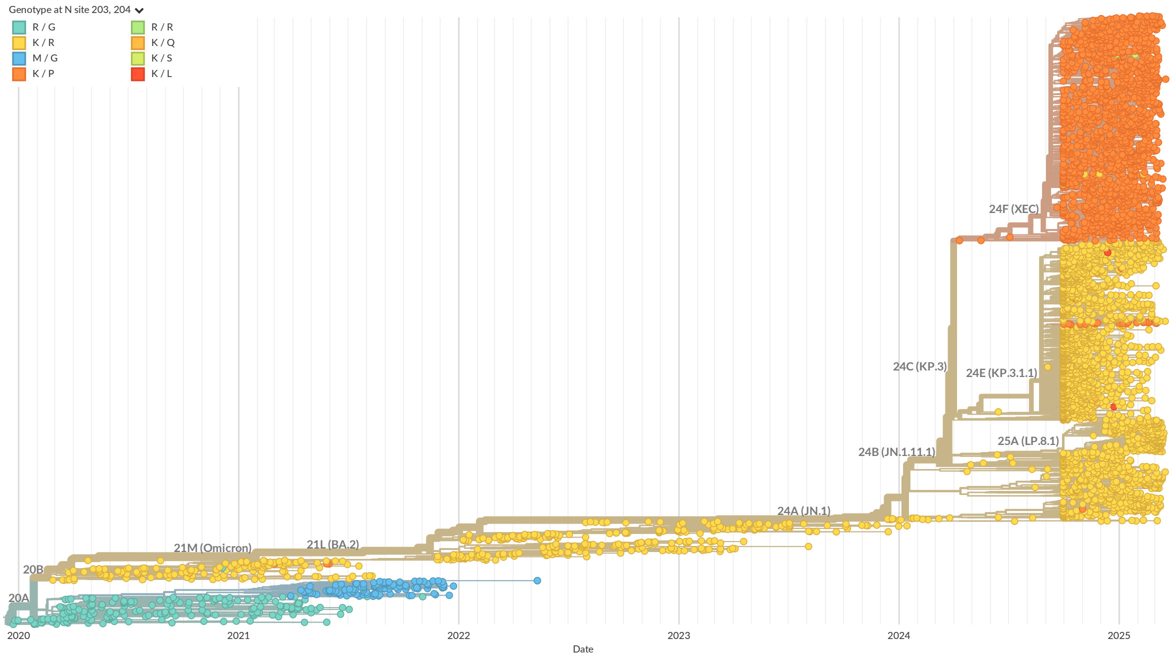
