## Supplemental Table S1 for "Evolution of a fuzzy ribonucleoprotein complex in viral assembly"

### Supplementary Table

#### Table S1. Peptide and Oligonucleotide Sequences

| designation | sequence |
|---|---|
| $N_{1\text{-}43}$ (N-arm) | MSDNGPQNQRNAPRITFGGPSDSTGSNQNGERSGARSKQRRPQ |
| $N_{1\text{-}43}$:P13L | MSDNGPQNQRNALRITFGGPSDSTGSNQNGERSGARSKQRRPQ |
| $N_{1\text{-}43}$:P13L,Δ31-33 | MSDNGPQNQRNALRITFGGPSDSTGSNQNG---GARSKQRRPQ |
| $N_{210\text{-}246}$ (LRS) | MAGNGGDAALALLLLDRLNQLESKMSGKGQQQQGQTV |
| $N_{210\text{-}246}$:G214C | MAGNCGDAALALLLLDRLNQLESKMSGKGQQQQGQTV |
| $N_{210\text{-}246}$:G215C | MAGNGCDAALALLLLDRLNQLESKMSGKGQQQQGQTV |
| $T_{10}$ | TTTTTTTTTT |
| SL7 | ACGUGGCUUUGGAGACUCCGUGGAGGAGGUCUUAUCAGAGGCACGU |
