## Supplemental Figure S2 for "Evolution of a fuzzy ribonucleoprotein complex in viral assembly"

**Supplementary Figure S2: Structural prediction of an RNP.** Top view (left) and side view (right) of an AlphaFold3 model of 5 N-protein dimers (ancestral) with 10 copies of SL7 RNA. One N-protein dimer is highlighted showing the two chains in blue and cyan. Three copies of SL7 adjacent to the highlighted dimer are shown in yellow, other copies are omitted for clarity. The top view shows the symmetrical arrangement of LRS helices forming a decameric core. The remainder of the linker, as well as the entire N-arm and C-arm are disordered and not meaningfully predicted. Due to significant disordered regions, the predicted model is not unique, and limited by the fact that multiple copies of SL7 are used as RNA ligand mimicking the biophysical RNP model.

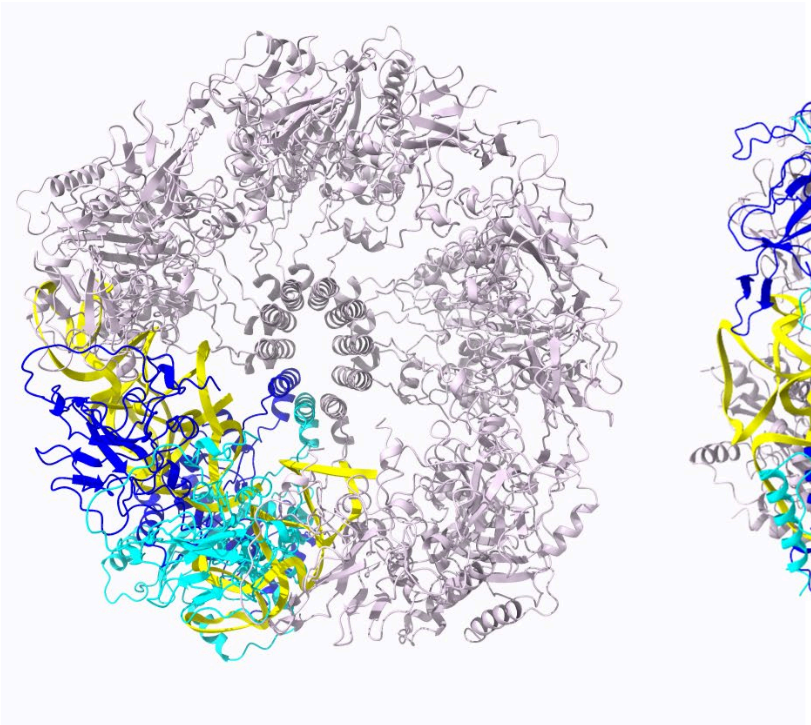

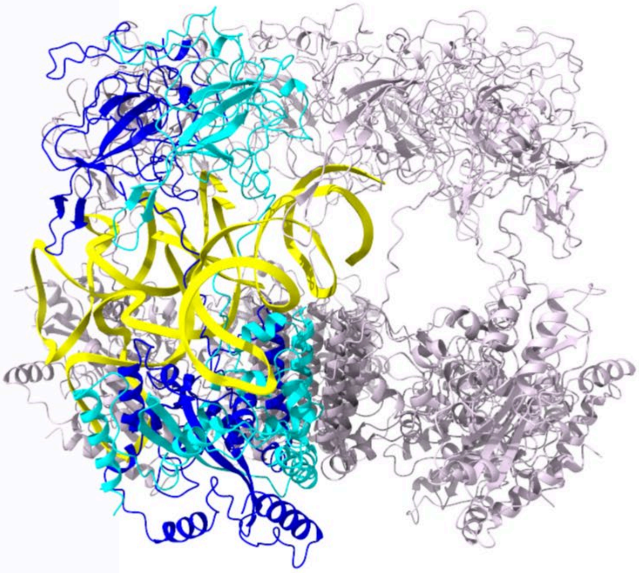
