## Supplemental Methods S1 for "Evolution of a fuzzy ribonucleoprotein complex in viral assembly"

### Supplementary Methods S1: Estimation of the Impact of Surface Binding on Solution Concentrations in MP.

Due to surface-binding of particles, their concentration in the solution above the surface will drop. How much of an effect is this under our experimental conditions?

Let a volume $V$ of sample at a concentration $c_0$ be applied to a well on the coverslip with area $A$, of which a fraction $f_A$ is observed in the image. If we count $n$ total surface binding events visible in the camera, assuming no unbinding, the total number of surface-bound molecules is $n/f_A$ and the number of molecules left in solution is $c_0 V N_A - n/f_A$. This means the concentration in the solution volume is now $c_1 = (c_0 V N_A - n/f_A)(V N_A)^{-1} = c_0 - n/(f_A V N_A)$, or a loss of $\Delta c = n/(f_A V N_A)$ occurred.

In our experiments, the values are: Sample volume $V$ = 10 µL with an initial concentration $c_0$ = 300 nM (N monomer), i.e., we start with ≈ $1.8\text{x}10^{12}$ molecules of N in solution. The well diameter is 3 mm, therefore the well area $A$ ≈ 7 $\text{mm}^2$, and the observed spot size is ≈46 $\mu\text{m}^2$, leading to the observed area fraction $f_A$ = $6.6\text{x}10^{-6}$ . Therefore, if we count $n$ = 10,000 particles (which exceeds the particle count in any of our experiments), the total number of particles absorbed would be $n/f_A$= $1.5\text{x}10^9$ (≈0.08% of the starting number), and accounting for (on average) approximately 10 copies of N per particle, this would amount to a reduction in concentration by approximately 0.8% or 2.5 nM. This is well within the concentration error.

This is in line with the observation that surface adsorption of proteins to glass is critical and needs to be prevented when working at picomolar concentrations (Zhao et al., 2014), but is ordinarily negligible when working at the mid nanomolar concentration range and non-functionalized surfaces.

It should be noted that this decrease in concentration is different from the decrease in count rate with time, which is governed by saturation of surface sites – an effect that diminishes the reduction of solute concentration above the surface with time. Finally, many particles exhibit desorption events, visible in a peak at negative contrast values, which will also further reduce the impact of surface adsorption on the solute concentration above the surface.
