## Supplemental Figure S1 for "Evolution of a fuzzy ribonucleoprotein complex in viral assembly"

**Supplementary Figure S1: Possible heterogeneity of RNPs from alternate configurations of N dimer subunits**. Only the N protein scaffold is shown for clarity. The nucleic acid binding domain at N-terminus (NTD) is indicated in blue, the dimerization domain (CTD) in green, and the L-rich region (206-247, LRS) folded into helices is shown as yellow rods. The other disordered regions are shown as black lines. (A) Based on the cartoon of **Figure 1C**, an RNPs is sketched where two of the 6 dimer subunits have flipped LRS orientations, resulting in the swapped position of NTD and CTD subunits. (B) Arrangement of 5 dimer subunits. (C) Arrangement of 7 dimer subunits. (D) $N_{210-419}$* constructs lacking the NTD are capable of forming RNPs of similar size, conceivably substituting NTD binding sites for nucleic acid by additional CTD subunits up-side-down orientation. (E) Mixed RNPs combining $N_{210-419}$* and full-length N protein different orientations.

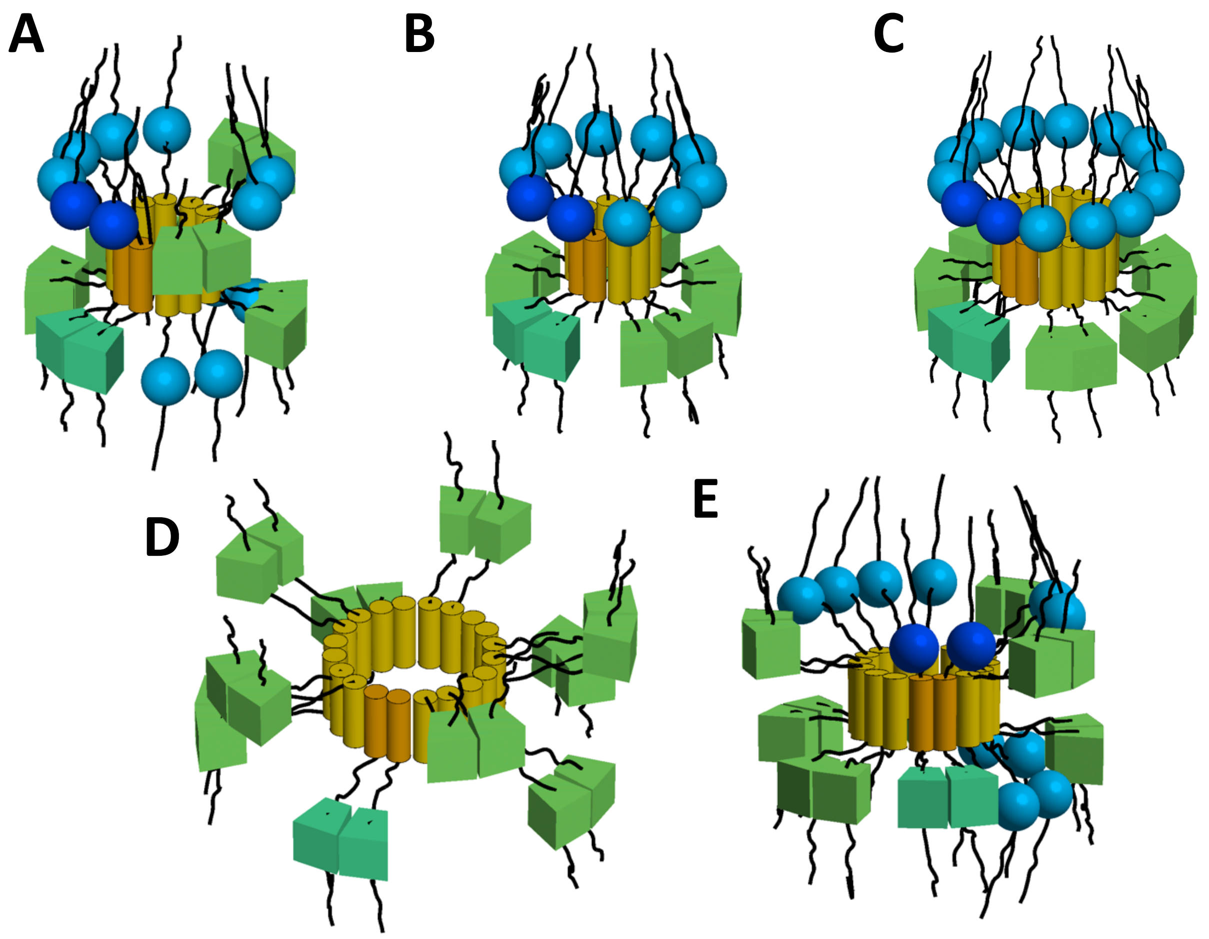
