## Supplemental Figure S3 for "Evolution of a fuzzy ribonucleoprotein complex in viral assembly"

**Supplementary Figure S3**: (A) Electron micrograph of negatively-stained N-arm peptide $N_{1-43}$:P13L after equilibration at 10 μM in 20 mM HEPES, 150 mM NaCl, pH 7.50. (B) As a control, electron micrograph of negatively-stained C-arm $N_{364-419}$ under the same conditions.

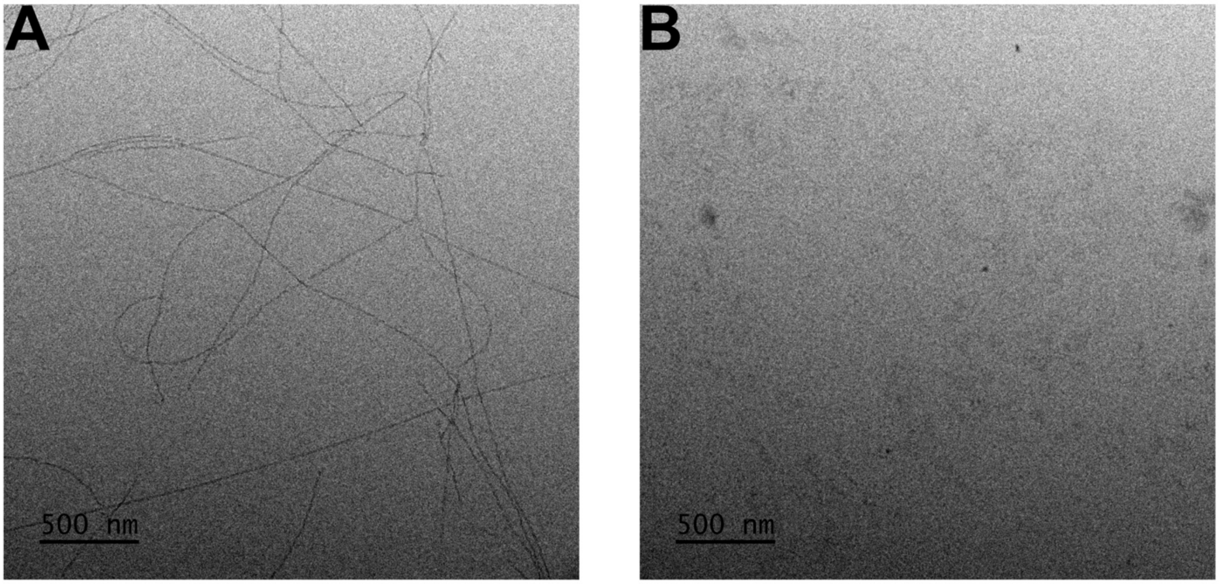
