## Supplemental Figure S4 for "Evolution of a fuzzy ribonucleoprotein complex in viral assembly"

**Supplementary Figure S4:** Comparison of WT and P13L structure predictions in ColabFold for a 12mer of the peptide $N_{10\text{-}20}$.

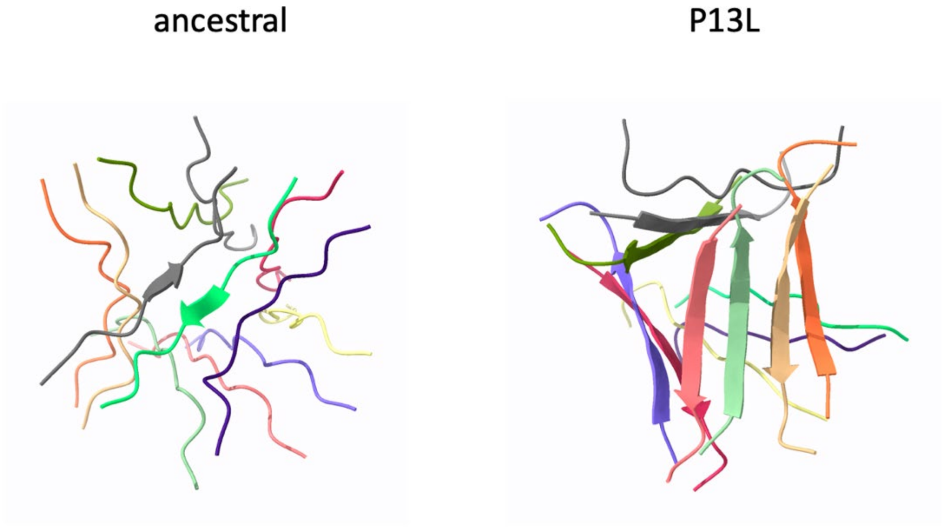
