## Supplemental Figure S5 for "Evolution of a fuzzy ribonucleoprotein complex in viral assembly"

**Supplementary Figure S5: Size distribution of G214C cysteine mutant LRS peptides.** Shown are peptides $N_{LRS, 210\text{-}246}$:G214C reduced (blue) vs unreduced (red). (A) Autocorrelation functions (circles) and best-fit size-distribution fits (solid lines). (B) Best-fit hydrodynamic radius distributions.

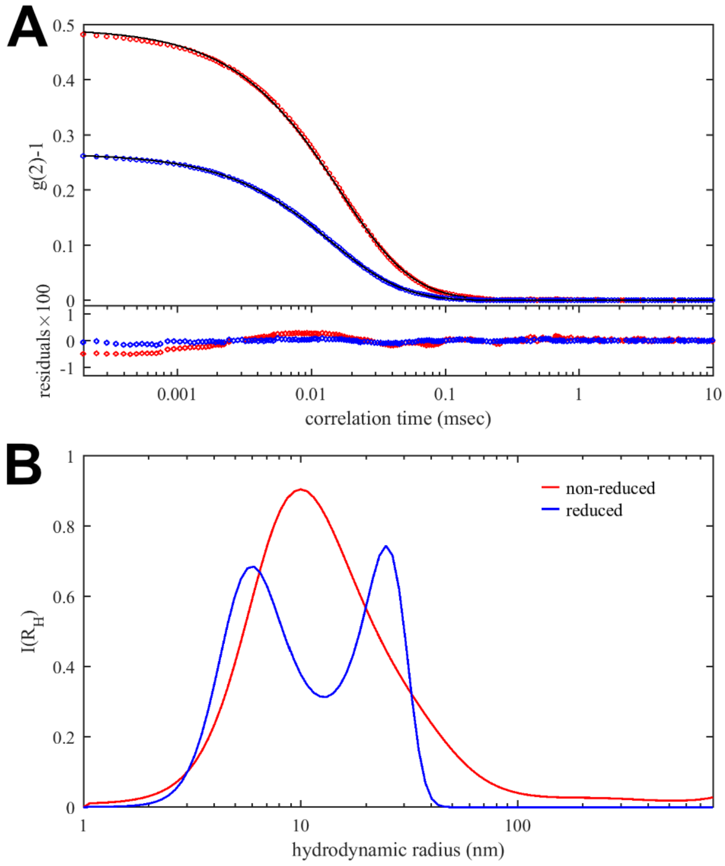
